# Glioblastoma disrupts cortical network activity at multiple spatial and temporal scales

**DOI:** 10.1101/2022.08.31.505988

**Authors:** Jochen Meyer, Kwanha Yu, Estefania Luna-Figueroa, Ben Deneen, Jeffrey Noebels

## Abstract

The emergence of glioblastoma in cortical tissue initiates early and persistent neural hyperexcitability with signs ranging from mild cognitive impairment to convulsive seizures. The influence of peritumoral synaptic density, growth dynamics, and spatial contours of excess glutamate upon higher order neuronal network modularity is unknown. We combined cellular and widefield imaging of calcium and glutamate fluorescent reporters in two GBM mouse models with distinct synaptic microenvironments and growth profiles. Functional metrics of neural ensembles are dysregulated during tumor invasion depending on the stage of malignant progression and tumor cell proximity. Neural activity is significantly elevated during periods of accelerated tumor growth. Abnormal glutamate accumulation precedes and outpaces the spatial extent of baseline neuronal calcium signaling, indicating these processes are uncoupled in tumor cortex. Distinctive excitability homeostasis patterns and functional connectivity of local and remote neuronal populations support the promise of precision genetic diagnosis and management of this devastating brain disease.

## Introduction

Glioblastoma (GBM), the most aggressive form of brain cancer, exploits the complex interplay between neurons and glia to create a microenvironment favorable for its own expansion yet hostile to network excitability homeostasis, impeding efforts to achieve long-term clinical remission. Nearly 60% of GBM cases present with seizures and develop pharmacoresistant tumor-related epilepsy (TRE), substantially reducing quality of life despite tumor resection ^1–3^. At the leading edge, a complex sequence of molecular and cellular remodeling leading to a hypersynaptic microenvironment ^4, 5^ with early loss and impairment of interneurons ^6–8^ promotes epileptogenesis and impairs cognitive processes. Distinguishing between healthy and epileptic tissue during tumor resection, as well as defining whether network hyperexcitability in areas more distant than 1 mm from the tumor margin is recruited and how this remodeling can be suppressed to regain healthy cortical function remain major treatment challenges.

Recent work has demonstrated that GBM progression can unfold nonlinearly over time^9^ and is intimately interconnected with TRE in a multifactorial fashion ^10–15^. GBM preferentially kills or dysregulates inhibitory interneurons ^16^ and modifies synaptogenesis ^4, 7, 11^ in a tumor driver gene-dependent manner ^17^. We previously showed in an immunocompetent murine model that GBM causes a gradual increase of electrographic cortical hyperexcitability coinciding with a reproducible pattern of peritumoral neuronal cell death, particularly in fast-spiking parvalbumin-positive (PV+) interneurons, degradation of perineuronal nets with widespread microglial activation and waves of spreading cortical depolarization that precede the onset of seizures, all providing evidence of this excitotoxicity ^7^. Similar pathology and GBM-induced changes in neurovascular coupling and neural activity dysregulation using human glioma cells implanted in mouse cortex offer additional evidence that improved understanding of malignant progression in diverse tumor subtypes will enable novel therapeutic opportunities ^8, 10, 12, 18^. With the advent of precision genetic profiling of human GBM ^19^, correlating the heterogeneity and pace of ambient pathophysiology with the tumor’s molecular profile may translate into improved individualized management of GBM cortical comorbidity.

GBM corrupts the synaptic microenvironment to facilitate its spread, and peritumoral glutamate is a well-established candidate mechanism underlying GBM hyperexcitability ^20–22^. Excess glutamate release is an expected feature of a hypersynaptic network, however the precise onset, temporal progression and spatial extent of neuron-tumor crosstalk relative to local and distant synaptic activity, and its variation among different tumor types, has never been determined. To address this variability, we compared a previously reported in utero electroporation (IUE) model of glioma (henceforth referred to as 3xCr)^4, 7, 17^ with tumors that included IUE addition of GPC6, a glypican family member of glial-secreted factors that promote synaptogenesis ^23^, since recent studies demonstrate that other glypican family members can promote glioma growth and TRE^17^. We combined chronic cortical imaging of peritumoral neural activity using calcium and glutamate reporters over a prolonged 3-month period of tumor expansion in awake mice. Simultaneously recorded EEG, locomotion, whisker and eye movements allowed us to control for the behavioral and attentional state of the animals during imaging. We analyzed peritumoral activity both mesoscopically at low spatial magnification/high temporal resolution covering both hemispheres, and at high magnification (2-photon excitation microscopy) in the same animals to pinpoint the distance of individual neurons from tumor cells at the leading edge. Our data reveal a nonlinear sequence of changes in functional network connectivity over time, a clear genetic dissociation between the speed of tumor invasion, and a strong correlation between accelerated tumor growth and neuronal hyperactivity. Spatially and temporally aberrant glutamate accumulation precedes and outpaces the spatial extent of neuronal calcium signaling, providing the first evidence that these processes are not tightly coupled.

## Methods

### Animals

All animal procedures were carried out under an animal protocol approved by the Baylor College of Medicine IACUC. Mice were housed at 22 deg C under a 12h/12h/ on/off light cycle. IUE-CRISPR mice were generated on CD-1 and C57BL/6 backgrounds as previously described ^4, 7, 17^ (see below). We used WT CD-1 mice for AAV-GCaMP, jrGeco1a, and iGlusnfr injections, or thy1-GCaMP6s-4.20 expressing mice on a mixed CD-1 x C57-BL6/J background.

### In-utero electroporation (IUE) model of GBM

In utero electroporation was performed on embryonic day 14.5 as described previously^24^. After pregnant moms were anesthetized, the uterus was exposed and a plasmid cocktail was injected into the lateral ventricles of developing embryos. Progenitor cells lining the lateral ventricles were electrotransfected, after which the uterus was reinserted, and surgical openings were closed. The injection mixture included a single construct which expressed previously reported guide sequences to target Pten, Trp53, and Nf1, along with the Cas9^4, 174, 15^. A piggyBAC transposable system was utilized to label cells with a GFP reporter and/or overexpressed GPC6.

### Immunohistofluorescence analysis

After IUE, mice were born and aged to post-natal day 30, at which point mice were euthanized and brains were dissected after transcardial perfusion of phosphate buffered saline (PBS) followed by 4% paraformaldehyde. After a brief post-fix, tissues were cryoprotected in 30% sucrose overnight. Brains were then embedded in Tissue-Tek OCT Compound (Sakura Finetek USA, 4583). Brains were sectioned at 40µm thickness and processed with antigen retrieval for their associated antibody staining. For BrdU-cell proliferation analysis, 2 hours before collection, 100 μg BrdU (in PBS) per gram body mass were delivered through intraperitoneal injection. Mouse brains were collected, frozen and sectioned as described above. Before blocking, sections were incubated in 2 N HCl at 37 °C for 30 min and neutralized with 3.8% sodium borate for 10 min at room temperature. For synaptic staining, sections were treated in 10mM sodium citrate, .05% Tween 20, pH 6.0 at 75°C for 10 minutes.

The following primary antibodies were used (at their associated dilutions, including catalog number (cn), and lot number(ln)): rat anti-BrdU (BU1/75 (ICR1), 1:200; abcam, cn: ab6326, ln: GR3269246-1), mouse anti-gephyrin (1:500; Synaptic Systems, cn: 147011, ln: 1-64), rabbit anti-GFP (1:1,000; ThermoFisher, A-11122), mouse anti-PSD95 (7E3-1B8, 1:500; ThermoFisher, cn: MA1-046, ln: LI147875), guinea-pig anti-VGAT (1:500; Synaptic Systems, cn: 131004, ln: 2-41), guinea-pig anti-VGLUT1 (1:2,000; Millipore, cn: AB5905, ln: 3193844). We used species-specific secondary antibodies tagged with Alexa Fluor 488, 568, or 647 (1:1,000, ThermoFisher) for immunofluorescence. After Hoechst nuclear counter staining (ThermoFisher, H3570, 1:50,000), coverslips were mounted with VECTASHIELD antifade mounting medium (Vector Laboratories, H-1000).

### Human Glioma Tissue analysis

Human glioma tissue microarrays were provided by Baylor College of Medicine’s Pathology and Histology Core. Sections were deparrafinized, rehydrated, and prepared for GPC6 staining (goat anti-mouse Gpc6, 10µg/mL, R&D Systems, cn: AF1053, ln:GJA0219031) through the following treatment: 3 × 3 min in xylene, 3 × 3 min in 100% ethanol, 3 × 3 min in 95% ethanol, 3 min in 80% ethanol, 5 min in 70% ethanol, 5 min in 50% ethanol, 5 min in ddH2O, antigen retrieval (10mM sodium citrated, .05% Tween 20, pH 6.0) at 95°C for 5 minutes, cool to room temp, 30 min in 3% hydrogen peroxide, 30 min in .1% Triton X-100 in PBS, and 30 min blocking with normal horse serum (provided in Vector Lab ImmPRESS HRP Horse Anti-Goat Polymer Detection Kit, MP-7405). After primary and secondary staining, signal was developed using the DAB Substrate Kit (Vector Lab, SK-1000) supplement with nickel. Sections were treated with Harris hematoxylin for 15 sec and the staining was blued with running tap water. Sections were then dehydrated and sealed with Permount before imaging.

### Survival analysis

After mice were born, they were observed for symptoms suggestive of tumors including (but not limited to) lethargy, hunched posture, decreased appetite, decreased grooming, trembling/shaking, squinting eyes, partial limb paralysis and abnormal gait, denoting the IACUC permitted endpoint. Statistical analysis was performed using the ggsurplot() function of survminer package (v 0.4.9)

### RNA-Sequencing and bioinformatics analysis

Samples from tumor brains were collected from endpoint, where tumor tissue was collected with the aid of fluorescence microscopy. Total RNA was isolated using the RNeasy Plus Mini Kit (Qiagen, 74134) according to the manufacturer’s protocol. Samples were prepared in biological replicates, n ≥ 3 per variant genotype. RNA integrity (RNA integrity number ≥ 8.0) was confirmed using the High Sensitivity RNA Analysis kit (AATI, DNF-472-0500) on a 12-Capillary Fragment Analyzer. Illumina sequencing libraries with 6-bp single indices were constructed from 1 μg total RNA using the TruSeq Stranded mRNA LT kit (Illumina, RS-122-2101). The resulting library was validated using the Standard Sensitivity NGS Fragment Analysis Kit (AATI, DNF-473-0500) on a 12-Capillary Fragment Analyzer. Equal concentrations (2 nM) of libraries were pooled and subjected to sequencing of approximately 20 million reads per sample using the Mid Output v2 kit (Illumina, FC-404-2001) on a Illumina NextSeq550 following the manufacturer’s instructions.

FASTQ files were quality controlled using fastQC (v0.10.1) and MultiQC (v0.9),11 and aligned to the mm10 reference genome via STAR (v2.5.0a).12 Count matrices and gene models were built from aligned files using Rsamtools (v2.0.0) and GenomicFeatures (v1.32.2) in R (v3.5.2). Using DESeq2 (v1.20.0) samples were normalized and analyzed for differential gene expression where GPC6 tumors were compared to 3xCr. Genes were considered significantly differentially expressed with a fold change > ±1.5 at P < .01. Ontology analysis was performed with the assistance of the Enrichr tool (maayanlab.cloud/Enrichr/).

Human expression and survival correlation analysis was performed utilizing the GEPIA online tool. Low and high expression cohorts were set at the bottom and top 50%, respectively. Log2 fold change cutoff was set at 1, P-value cutoff was set at .01, and match normal data were set to TCGA normal and GTEx data.

### EEG and headpost implant, virus injection, craniotomy

Mice were induced with 3-4% isoflurane in oxygen at 2 L/min flow rate until breathing slowed to ∼60 bps. Deep anesthesia was confirmed via tail pinch. Hair was removed by electric clippers (Wahl, Sterling, IL, USA), and depilatory cream (Nair, Church & Dwight, Ewing, NJ, USA). The mouse was immobilized in a stereotactic frame with a gas-anesthesia adapter for neonatal rats and blunt-tip ear bars aimed at the upper mandible joint (Kopf instruments, Tujunga, CA, USA). The scalp was disinfected using 3 alternating applications of betadine scrub and 70% EtOH, and the animal was covered with a fenestrated sterile drape sparing only the disinfected scalp. All surgical instruments were autoclaved prior to use. Using pointed tip surgical scissors (WPI, Sarasota, FL, USA), a ∼1×1 cm square skin flap was removed to expose the skull bone. Using a no. 10 scalpel blade and serrated forceps, the periosteum was removed, and residual bleeding was absorbed with sterile cotton swabs. A thin layer of Vetbond cyanoacrylate (3M, Maplewood, MN, USA) was applied to close the scalp wound. Next, a custom-made EEG adapter with attached wires (2 recording channels, one reference, each made from Teflon-coated silver (WPI, Sarasota, FL, USA) wire with ∼0.5mm tips exposed, beveled using 600 grit sandpaper and chlorided in NaClO) was attached using Krazy glue (Elmer’s, Westerville, OH, USA) to the skull bone ∼3 mm anterior of lambda. A dental drill with ¼ carbide burr tip (Midwestern, Henry Schein, Melville, NY, USA) was used to make 3 burr holes, one over the cerebellum (bregma -3 mm, 2.5 mm lateral), and two at lambda +1 mm, 5 mm lateral on both hemispheres, making sure not to break through the bone but to create small cracks exposing the underlying dura mater. The silver wire tips were carefully inserted just below the bone to touch the dura, and the holes were immediately sealed with a small drop of Vetbond. A mixture of dental cement powder and charcoal powder (10:1) was mixed with dental cement fluid (Lang dental, Wheeling,IL, USA), and applied to the skull in a donut shape, approx. 5 mm tall and creating a ∼15 mm diameter well. The custom-designed aluminum head bar (emachineshop, Mahwah, NJ, USA) was gently pushed into the dental cement to sit flush with the skull. A small amount of Vetbond was added to the inside walls of the headpost to promote permanent attachment. After the dental cement cured, excess cement was removed using the dental drill. A nanoliter injector (Nanoject II, Drummond scientific, Broomall, USA), was used to inject AAV solutions. Glass pipettes were pulled using a Sutter P-87 horizontal pipette puller (Sutter instruments, Novato, CA, USA) and tips were broken on the filament of a vertical puller (Narishige, Japan) to create a sharp, ∼10-20 µm wide opening. The pipette was backfilled with corn oil, and ∼5 µL virus solution was aspirated from a sterile piece of parafilm using the Nanoject II. Depending on the experimental paradigm, we used one of the following (combinations of) AAV’s:

pGP-AAV-syn-jGCaMP7f-WPRE (Addgene, Watertown, USA), diluted 1:2.5 in sterile Cortex buffer (CB),

or AAV-FLEX-GCaMP8m + AAV-CamKIIa-0.4-Cre, diluted 1:1.5 in sterile CB

or AAV-jrGeco1a, diluted 1:3 with sterile CB,

or AAV-FLEX-iGlusnfr-184 + AAV-Ef1α-Cre, diluted 1:1.5 in sterile CB,

or AAV-FLEX-iGlusnfr-184 + AAV-CamKIIa-0.4-Cre, diluted 1:1.5 in sterile CB.

A total of 8 or 6 (depending on the unobstructed skull area available) equidistant injection locations were selected with the goal to maximize coverage of sensory and posterior motor cortex. For each hemisphere, these were at [bregma –3.5 mm; 2.5 mm lateral], [bregma –2 mm; 1 mm lateral], [bregma –2 mm; 4 mm lateral], and [bregma –0.5 mm; 2.5 mm lateral]. At each location, a total of 400 nL AAV-solution at 2 depths, ∼300 µm, ∼600 µm, were injected into the cortex in 9.2 nL/pulse increments separated by 10s. After the last pulse in each location, the pipette remained in place for 5 min to minimize backflow and promote virus diffusion. The Nanoject was mounted at a 25-degree angle relative to the skull surface at each location and actuated by a manual and 1-direction motorized micromanipulator (WPI, Sarasota, FL, USA) at speeds of ∼600 µm per min. After the last injection, the skull was covered with Vetbond and dental cement. Mice were allowed to recover for ∼10 days post injection with meloxicam analgesic administered for the first 3 days. For the second surgery, induction and stereotactic positioning were performed as described above. The dental cement covering the skull was removed. A single-pane #1 cover glass (22×11 mm, Thomas Scientific, Swedesboro, NJ) was cut and adjusted to the individual dimensions (approx. 7.5×6 mm) of the exposed skull bone by scoring the glass with a sharp-tip drill bit. The cover glass was then submerged in 70% EtOH for >10 min. The outline of the cut glass was transferred, and a corresponding groove carefully drilled while cooling the bone every 10-20s with compressed air and stopping short of the dura to avoid damage. A thin strip of bone covering the superior sagittal sinus was thinned but not removed. Before carefully removing the bone, it was soaked to soften for 5 min with sterile CB. Any superficial bleeding was controlled with small pieces of sterile surgifoam (Ethicon, Cincinnati, OH, USA), previously soaked in sterile CB. The dura was carefully removed by grasping, stretching and gently piercing it with a 30G needle, then peeling it off with serrated forceps. The cover glass was carefully placed and initially attached with Vetbond while gently applying downward pressure, ensuring that any gaps between the bone and the glass were filled with Vetbond without reaching the brain. After polymerization of the Vetbond, a few drops of cyanoacrylate glue were added to secure the glass in place, and finally a thin layer of dental cement around the edge of the glass was applied to seal it in place. Typically, this preparation allowed for >6 weeks of cellular resolution 2-p imaging and >10 weeks widefield 1-p imaging. If redness or signs of inflammation was seen, dexamethasone (0.5 mg/kg) was given for 3 days.

### In vivo 2-photon and 1-p widefield imaging

Mice were typically imaged every 4-6 days starting between P45 and P55, and every 1-3 days once spontaneous seizures were detected until the animal appeared moribund and was euthanized with CO_2_. (Avg time span between first and last imaging session was 48 ± 31 (SD) days). The mouse was headfixed and allowed to freely run on top of a custom-built Styrofoam wheel (12 cm diameter), whose rotational speed was recorded by an angular velocity encoder. Both imaging modalities are integrated in a Prairie Ultima IV (Bruker, Billerica, MA, USA) modified in vivo 2-photon microscope. One-photon image sequences were acquired with a pco.Edge 4.2 sCMOS camera (pco, Germany), mounted on top of the scan head of the Bruker Ultima IV microscope with a 2x objective lens (CFI Plan Apo Lambda 2X, Nikon, Melville, USA), yielding a raw (unbinned) resolution of 3.62 µm/pixel (FOV = 2060×2048 pixels). Widefield movies were acquired using 2x or 4x software binning at 100Hz (10ms exposure duration). Blue illumination light (for GCaMP indicators or GFP expressed in tumors) was generated by a Xenon arc lamp (Zeiss, Germany), guided through a 480/20 excitation filter. For tagBFP labeled tumors, a 400/40 excitation and 480/40 emission filter was used. For GCaMP emission and jrGeco1a or RFP excitation, a 525/70 filter was used, and for jrGeco1a and RFP emission, a 620/60 filter was used. For iGlusnfr (Venus) imaging, a 517/20 excitation and 554/23 emission filter set was used. Images were acquired at 100 Hz with 2×2 or 4×4 digital binning (1020×1024 or 500×512 pixel FOV, respectively) on a Lenovo P920 workstation (Lenovo, Morrisville, USA). EEG signals were amplified (HP 0.1 Hz, LP 5 kHz, gain x100) using a Model 1700 differential AC amplifier (A-M systems, Sequim, USA), digitized with a USB-6211 multifunction I/O device (National Instruments, Austin, USA) and recorded using WinEDR freeware (Strathclyde University, United Kingdom) at 10 kHz. Pupil size of the right eye was detected with IR LED illumination and recorded at 30 Hz with a GC660 IR camera (Allied Vision Technologies, Newburyport, USA) using custom Matlab routines (Mathworks, Natick, USA). The position and movement of the animal’s head, whiskers, front paws and frontal parts of the body was monitored and recorded at 30 Hz with another IR camera triggered by the WinEDR output signal generator (model SC1280G12N, Thorlabs, Austin, USA). These recordings were used to detect and exclude periods containing movement artifact that might contaminate the pixel by pixel analysis of imaging data.

**2-photon images** of calcium and glutamate reporter activity were acquired with a Prairie Ultima IV 2-photon microscope using a ×25 objective, 1.1 NA, or a ×16 objective, 0.8 NA, at 920 nm (GCaMP7/8) or 1000 nm (jrGeco1a) under spiral (10–20 Hz frame rate) or resonant scan mode (30-35 Hz). We used a 525/70 nm emission filter for GCaMP6/7/8 indicators, a 554/23 nm filter for iGlusnfr, and a 620/60 nm filter for jrGeco1a emission. We targeted primarily L2/3 (mean depth below pia: 163 μm, range 100–240 μm). We also imaged some FOVs in L4 (mean depth below pia: 383 μm, range 360–395 μm), and several FOVs in L5 (mean depth below pia: 570 μm, range 510–640 μm). Laser output power under the objective was kept below 50 mW, corresponding to ∼20% of power levels shown to induce lasting histological damage in awake mice ^25^. Mice were imaged while awake, head-posted in a holding frame and allowed to run freely on a circular treadmill.

### 1-photon image preprocessing

Widefield images were processed using a custom-written Matlab pipeline partially based on previously published analysis methods [suite2p/caiman/seqnmf/mesoscale brain explorer/ofamm toolboxes ^26–28]^. Briefly, recordings were saved by the pco acquisition software as multiple 2GB tiff stacks. These were loaded into MATLAB, downsampled to a spatial resolution of ∼36 µm/pix, motion-corrected ([normcorre-function], Matlab ^29^), multiplied by a mask tiff file to suppress any artifactual signals from the vasculature (manually constructed in imageJ), and converted into ΔF/F movies saved as a single .h5 file. The other acquired voltage traces corresponding to wheel motion, visual stimulus, IR camera trigger, and EEG were then downsampled and aligned to the ΔF/F traces.

Snapshots (average of 20 frames) of baseline activity during quiet wakefulness were taken at full resolution (2048×2048 pix) and 100 ms exposure time. To resolve the magnitude of progressive change in neuronal activity and glutamate levels during these intervals, we quantified the evolution of baseline peritumoral calcium signaling and glutamate intensity in mice at a repeated sampling time interval ranging from 3-7 days over a period of 6-11 weeks (n=8 mice with calcium indicator, n=7 mice with glutamate indicator), as determined by survival. To assess regional changes at sequential time intervals, we calculated the coefficient of variation (CV) for each pixel between consecutive imaging time points for calcium, glutamate and tumor indicators by first scaling all images from each animal equally, then calculating the ratio of images taken at consecutive recordings, calculating the standard deviation across of all pixels, dividing by the number of days between the sessions, and normalizing by the mean intensity of all pixels of the resulting ratio image (see figures 3, 4). The colorized panels show a relative increase in tumor CV/day in red (up to +50%) and a decrease in blue (up to -50%, white = no change). To summarize the overall progression, we binned the recordings into time intervals of 10 days between P45 and P135 and computed the 90^th^ percentile of the resulting distributions for each bin to account for potential sampling bias towards more frequently sampled animals and potential outliers.

### Analysis of widefield calcium signal in relation to tumor growth

We used mesoscopic one-photon calcium imaging of bilateral cortical FOV’s spanning on average ∼45 mm^2^ at 100 Hz frame rate in 4 3xCR tumor animals. Tumor coverage of the FOV was measured for each recording, and tumor growth rates were calculated in µm^2^/day. Based on our observation that tumors generally either existed in a prolonged state of slow growth (<10^5^ µm^2^/day) or enter an accelerated growth rate of >10^5^ µm^2^/day (figure 3D), we divided the neural recordings into slow growth (n = 20 recordings from unique time points) or fast growth (n = 13 recordings) categories. Detrended and normalized ΔF/F image sequences were spatially downsampled by a factor of 8, resulting in a final resolution of ∼240 microns per pixel. Active running and whisking episodes were computed and excluded to select images corresponding to 300 sec of quiet wakefulness for each recording (suppl fig 1a). For each pixel, a ΔF/F trace was extracted, and calcium events were identified using a thresholding algorithm, taking into account variations in the baseline noise level for each pixel. From here, we defined the following metrics that provide information about different aspects of the calcium activity patterns observed (suppl fig 1c).: 1) “activity per min”: the mean of the ΔF/F trace computed over the 300 sec trace. 2) “events per sec”: the rate of identified calcium transients after thresholding at 3SD above the median, divided by 300 sec. 3) “mean amplitude”: the maximum ΔF/F value of each calcium event averaged over all identified events per pixel. This corresponds to a measure of the maximal firing rate achieved during a single calcium transient 4) “mean area”: The area under each event transient was averaged over the 300 sec trace for each pixel. This corresponds to a relative measure of sustained or “integrated” firing rate during the duration of a calcium transient. Note that this measure does not distinguish between a long, low-amplitude event and a short, high-amplitude event of the same area under the curve. 5) “rhythmicity”: This metric was computed by generating a distribution of all inter-event intervals and calculating the inverse of its standard deviation divided by the mean. Note that this metric shows the regularity of calcium events but does not contain information about the frequency at which they occur. Changes in these metrics for all FOV pixels pooled together over time were quantified using a non-parametric t-test equivalent (Wilcoxon ranksum test with multiple comparison correction (“WR/mc”) across recordings. For each recording session, a reference image of the tumor (using either a green/red filter for RFP, blue/green for GFP, or 400 nm/blue for BFP) was acquired, spatially downsampled and aligned with the calcium signal image to obtain a pixel mask of the tumor mass location. Tumor growth rate for each period between recording sessions was calculated by dividing the difference in tumor fluorescence between consecutive recordings by the number of days in between. Tumor spatial coordinates were extracted, and the distance between each pixel and the nearest part of the tumor edge was computed (suppl. fig 1b). A linear regression between all tumor distances and the 5 metric values was performed and adjusted R^2^ as a goodness-of-fit parameter calculated. The tumor distances were circularly shuffled 500,000 times, and the regression was performed for these bootstrapped repetitions (suppl. fig 1d). Significance (p-value) was then computed as the fraction of bootstrapped trials with a higher R^2^ than the actual value. Significance was quantified as a function of tumor growth rate. For recordings with significant changes, FOV pixels were then divided into distance bands at 0.75-mm steps from the tumor, and distance-related changes were depicted as shaded errorbar plots showing the defining features of the metric distributions in each band.

### Analysis of 2-photon cellular calcium signals and network activity

Raw tiff image stacks were loaded into matlab, motion corrected using normcore, and CNMF was used to identify active neurons in the FOV. ΔF/F traces were computed for each neuron, followed by deconvolution and neuropil contamination correction (adapted from suite2p-wrapperdeconv function ^26^, yielding a deconvolved ΔF/F matrix (termed “dΔF/F”) that was used for all subsequent analysis steps. Next, whisking, wheel velocity, EEG and photodiode voltage traces were downsampled and aligned to the dΔF/F traces. Periods of active whisking and running were excluded for the remainder of the analysis. Deconvolved traces were then thresholded by identifying the bottom 20^th^ percentile of all data (corresponding to random noise), and its median and SD. After thresholding above median+3SD, individual deconvolved calcium transients were identified, and 4 metrics were extracted: 1) overall (summed) dΔF/F per min, 2) events/sec, 3) amplitude of the transients, 3) duration (or length) of the transients. The 4^th^ metric we computed for each neuron was the clustering coefficient (CC): To identify coactive clusters of neurons comprised of >2 neurons, the clusterONE algorithm (BrainConnectivityToolbox, matlab ^30^) uses the previously computed pair-wise Person correlation coefficient matrix as its weighted network input, and identifies potentially overlapping groups of at least 3 associated neurons. The CC of each neuron represents the tendency of its neighbors to be interconnected (see equation (9) in ^31^).

We acquired these neuronal activity measures from FOV’s at different locations (inside and outside the tumor margin), at different time points (early (P40-49), mid (P-50-59) and late (P60-100) stage), and across tumor genotypes (3xCR, GPC6 and non-tumor control). To analyze chronic changes in activity patterns, we split up the GPC6 and 3xCR recordings into 3 time bins: early (P40-49, mean 43.6 +/- 0.84 sem), mid (P50-59, mean 54.9 +/- 1 sem), and late (P60-100, mean 89.2 +/- 4.3 sem) in order to compare across 3 dimensions: time, distance from the tumor, and tumor genotype. The control late group spanned P80-P129. For the comparison between areas inside the tumor margin and tumor cell free surrounding areas, we included data only from mid-to late time bins, as we could not ensure that earlier time points left enough time to uniformly express the calcium indicator in all AAV injected animals. For each animal, we chose FOV’s that were either located within the infiltrating tumor margin, i.e. neurons were visibly intermixed with tumor cells (0-1mm from the solid (neuron-free) tumor core), or located within a ‘leading edge’ band of 1-2mm from the tumor margin, where there were no tumor cells visible inside the FOV We verified the lack of tumor cells in these leading edge FOVs by acquiring z-stacks, but could not rule out the presence of tumor cells beyond a depth of >600µm due to imaging limitations. Therefore, analogous statistical comparisons were performed independently across these 3 dimensions: First, we randomly chose equal numbers of neurons from each recording to ensure equal weighting of animals in each group, based on the FOV in each group with the lowest number of identified neurons (3xCR early: n=100, 4 animals, 3xCR mid: n=100, 5 animals, 3xCR late: n=100, 5 animals, 3xCR inside: 30, 5 animals; GPC6 early: n=50, 5 animals, GPC6 mid: n=100, 5 animals, GPC6 late: n=100, 5 animals, GPC6 inside: n=25, 5 animals ; control mid: n=100, 4 animals, control late: n=100, 5 animals). We studied a total of 11 3xCR animals, 9 GPC6 animals, and 8 control animals, and unique datasets from some animals were included in several groups. For each comparison between groups, pooled data distributions, which were generally not normally distributed (see violin plots in figures 6, 7), were compared using the nonparametric Wilcoxon ranksum test, and for each group the significance threshold was set at α = 0.05/n (n=number of comparisons) to account for multiple comparisons (termed “WR/mc test”). Percent change between compared groups was displayed with bar plots if deemed significant.

## Results

### Tumors with and without GPC6 generate distinct cortical infiltration dynamics

Our previous studies demonstrated how tumor intrinsic factors influence the local neighboring neurons and the larger synaptic network at critical stages of tumor hypeexcitability^4, 7, 17, 32, 33^. In an effort to gain more granular insights into these dynamics, we developed a multi-metric, longitudinal imaging assay system for prolonged, regular monitoring of tumorigenesis as shown in Fig 1.

**Figure 1:**
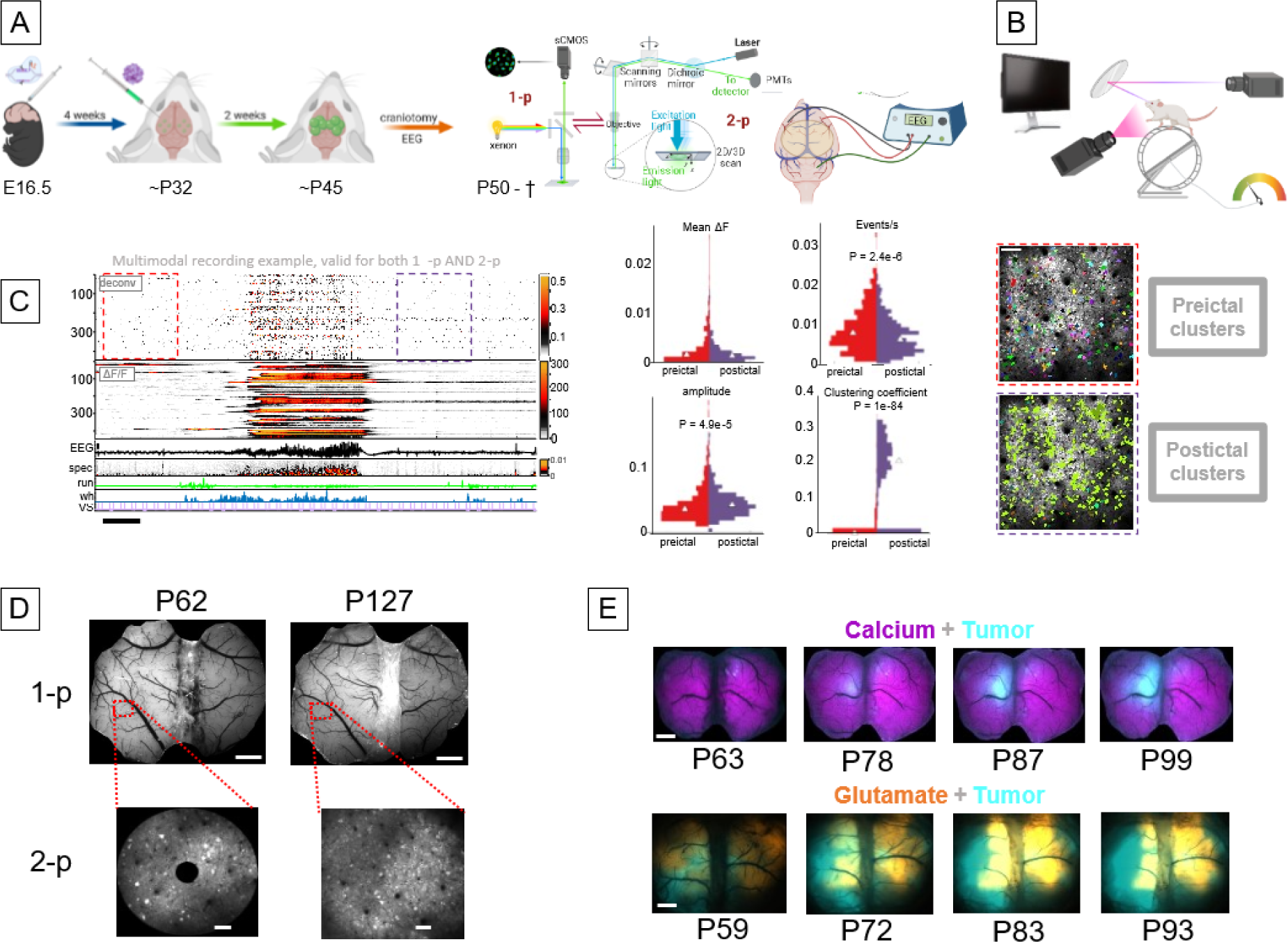
Integration of an IUE murine GBM model with multimodal recordings of cellular, widefield fluorescence and behavioral parameters. A) Schematic illustration of experimental design. IUEs are performed on E14.5 mouse embryos. At 1 month of age, AAV viruses are injected intracortically. After 2 weeks, a bilateral ∼7×5 mm cranial window and 2-channel EEG electrodes were installed on the mouse’s cranium. B) Schematic illustration of multimodal monitoring of the animal’s behavioral state: Running wheel velocity, pupil and eye movements, and whisking/frontal body movements were recorded. An LCD monitor was used to display visual stimuli. C) Time series raster plot of representative 2-photon imaging recording showing baseline (red dotted box), seizure, and postictal activity (black dotted box). Traces show ΔF/F calcium activity, EEG voltage, and mouse behavior illustrated in (B). Scale = 10 sec. Violin plots display distributions of deconvolved activity metrics extracted from the baseline (red dashed rectangle), and postictal (purple dashed rectangle) periods. White triangles = median. Preictal and postictal clusters shown to the right were computed from the same recordings. Neurons belonging to the same cluster share the same color. Scale = 0.1mm

To assess the potential contributions of GPC6 in gliomagenesis, we first mined its expression in human GBM. Utilizing established, publicly available datasets, we observed that GPC6 expression is significantly increased in both low grade glioma (LGG) and GBM compared to normal brain tissue (Fig 2A). To evaluate whether GPC6 could be functionally relevant to gliomagenesis, we compared the survival of human patients with differential GPC6 expression, finding that high GPC6 expressing patients demonstrated significantly worse survival (Fig 2B). Lastly, to validate the presence of GPC6 in human patients, we tested for GPC6 protein in a glioma tissue microarray, finding enhanced expression in tumor samples compared to normal brain (Fig 2C). Together, these findings suggest that GPC6 could be functionally relevant toward promoting gliomagenesis.

**Figure 2:**
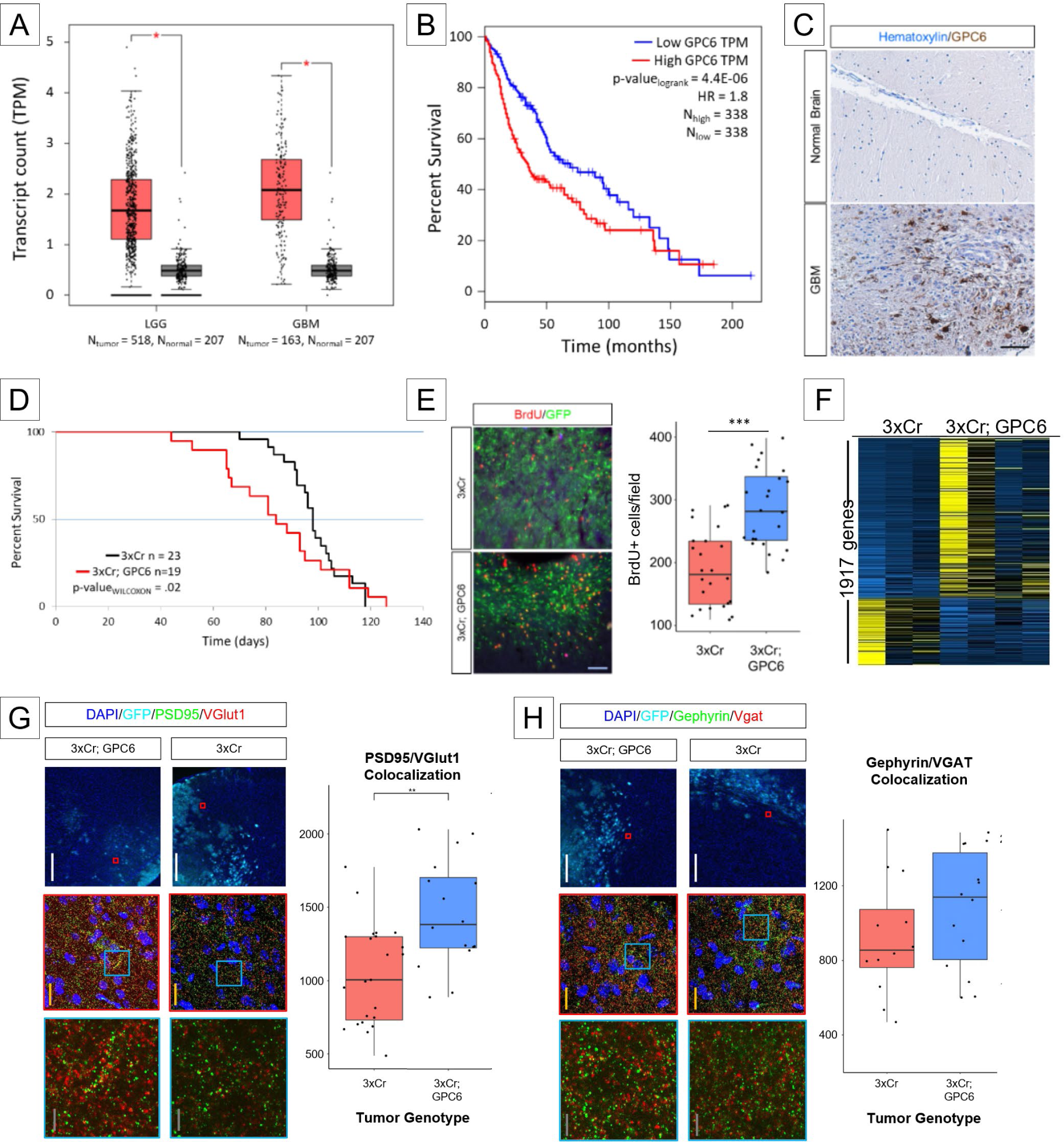
GPC6 is enriched in human and murine GBM. A) GPC6 expression analysis based on transcript count per million (TPM) from human datasets comparing low grade glioma (LGG), GBM, to normal non-tumor brain. Asterisks signifies p-values < .01. B) Kaplan-Meier survival analysis of human patient cohorts of LGG and GBM patients. Low and high TPM cutoff were set at below and above the 50^th^ percentile, respectively. HR – hazard ratio. C) Immunohistological staining for GPC6 on human GBM tissue (below) along with normal brain control (above). Scale bar = 100µm. D) Kaplan-Meier survival analysis comparing 3xCr (black) and GPC6 (red0 tumor bearing mice. p-values calculated through log-rank method. E) Representative immunohistofluorescence of BrdU incorporation (red) in tumor sections (green). Quantification of BrdU+ cells per field (100,000 µm^2^) p-values calculated by student t-test. *** < .001. N_biological_ = 4 brains. N_technical_ = 6 images per brain. F) Heatmap of differentially expressed genes between 3xCr and GPC6 tumors brains. Each column represents a single brain. First 3 columns are 3xCr brains. Last 4 columns are GPC6 tumors. Yellow – high expression. Blue – low expression. G) Representative images from immunohistofluorescence images of PSD95 (green) and vGlut1 (red) staining around tumor margins (cyan). Quantifications of colocalization of Psd95 and vGlut1. p-value calculated by student t-test. ** < .01. White scale bar = 300µm; yellow scale bar = 20µm; grey scale bar = 5µm. N_biological_ = 4 brains. N_technical_ = 6 images per brain. H) Representative images from immunohistofluorescence images of Gephyrin (green) and Vgat (red) staining around tumor margins (cyan). Quantifications of colocalization of Gephyrin and Vgat. p-value calculated by student t-test. No significant difference. White scale bar = 300µm; yellow scale bar = 20µm; grey scale bar = 5µm. N_biological_ = 4 brains. N_technical_ = 6 images per brain.

To test for this, we overexpressed GPC6 in our IUE mouse glioma model (Fig 2D). We observed that adding GPC6 significantly accelerated tumor associated death with a difference in median survival of 2 weeks (median survival 3xCr vs GPC6: 97 vs 83 days, respectively), while overall survival was similar between the two. Interestingly, this mimics the patterns observed in human survival studies. In concordance with the quickening of tumor burden, we observed a significant increase in BrdU incorporation (Fig 2E), suggestive of accelerated tumor growth.

To gain a molecular perspective on the changes GPC6 elicits on a tumor brain, we performed bulk tissue RNA-Sequencing (RNA-Seq) from endpoint 3xCr and GPC6 tumors and identified 1917 differentially expressed genes (p-value < .05, Fig 2F). Of these 1917, 1350 were upregulated in GPC6 tumor brains. Gene ontology analysis (Supp Fig 2) across different ontology databases revealed that the top categories were associated with synapses, neurons, or seizure/hyperexcitability. To assess changes in the neighboring synaptic microenvironment, we stained peritumoral sections for excitatory and inhibitory synapses in developing tumors at P30 (Fig 2G-H) and found a significant increase in excitatory synapses but no significant change in inhibitory synapses in the presence of GPC6.

We analyzed the spatiotemporal patterns of cortical tumor invasion, comparing the 3xCR tumor model with and without the overexpression of GPC6. First, we calculated the CV/day (i. e. coefficient of variation of pixel-wise daily rate of intensity fluctuation) between serial tumor fluorescence samples (see methods, Fig 3A,B). In contrast with the early large dynamic changes in tumor fluorescence in the 3xCR model, the GPC6 animal (Fig 3B) exhibits significantly attenuated variations in tumor fluorescence CV over time (images calculated from 7 time points between P62 and P114). We analyzed a total of 11 3xCR and 8 GPC6 animals and found that the CV/day was on average 49% lower for GPC6 than for 3xCR tumors (0.032 +/-0.005 sem vs 0.065 +/- 0.011 sem, p = 0.032 WR test, figure 3C). In addition, quantification of tumor coverage rates (in μm^2^/day) as a function of time revealed early elevation of GPC6 infiltration, and sustained growth rates of 3xCR tumors that were lower at late stages in GPC6 animals (3D). In each recording session we verified the presence of tumor cells and the outline of the tumor margin by acquiring cellular-resolution 2-photon z-stacks (Fig 3E). These results reveal that 3xCR tumors relentlessly infiltrate cortical territory throughout disease progression, whereas GPC6 tumors first infiltrate aggressively, but grow less dramatically later once they reach cortical layers 1-4. This unexpected non-uniform growth rate and laminar microenvironment effect would not have been made without intensive chronic in vivo imaging.

**Figure 3:**
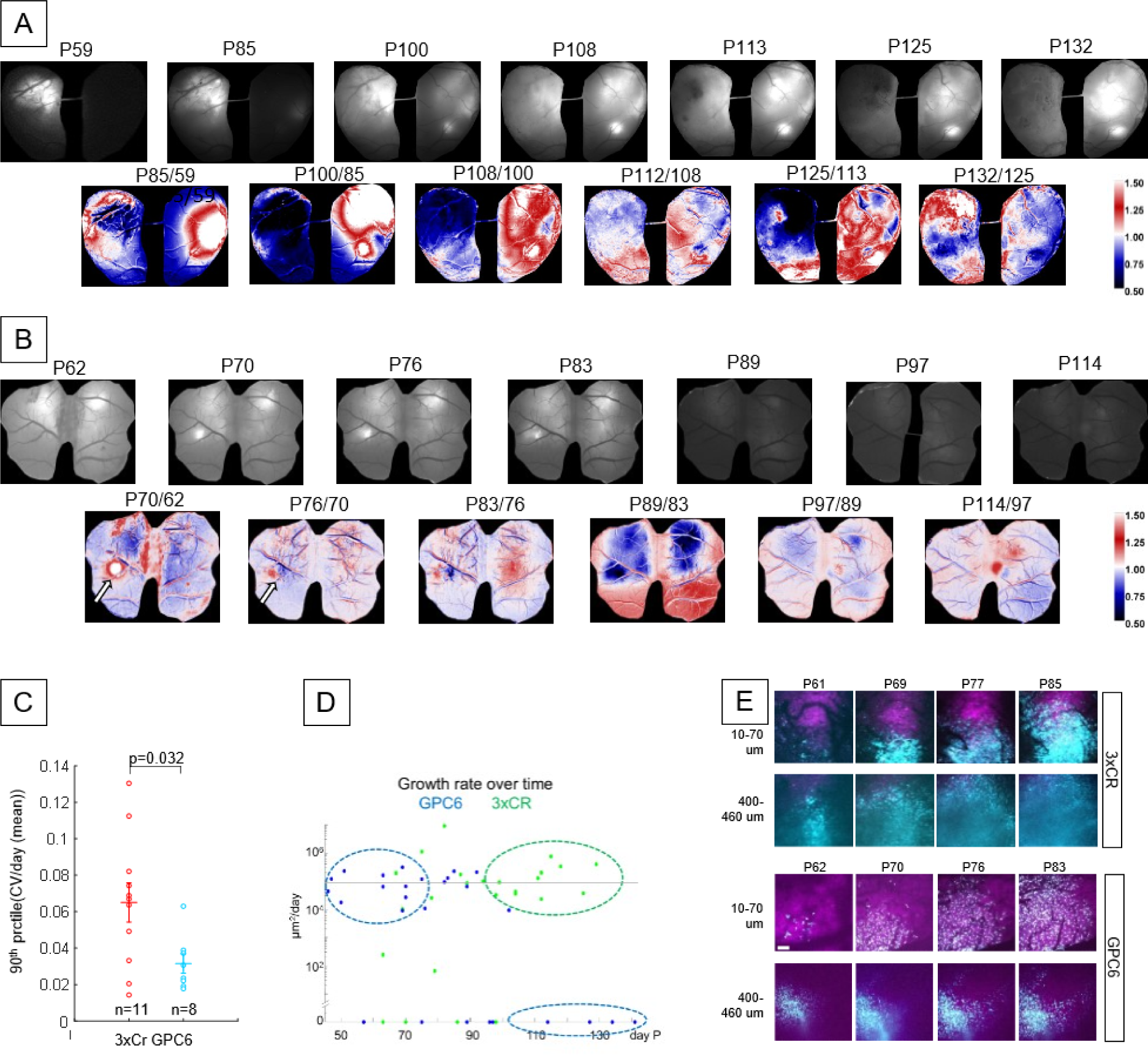
GPC6 induces early tumor growth. A) Top row: representative mesoscopic images of tumor labeled fluorescence over time of 3xCr tumor brains. Age of mice in postnatal days listed above image. Bottom row: colored panels, generated from above monochrome intensity, colored based on changes to signal strength at location between 2 time points. B) Analogous analysis to (A) for GPC6tumor animal. C) Quantification of tumor images from 11 3xCR and 8 GPC6 comparing 90^th^ percentile data point of CV/day. Mean 3xCr = 0.065+/-0.011 sem. Mean GPC6 = 0.032+/-0.005 sem, p=0.032, Wilcoxon ranksum test). D) Mapping tumor growth rate by age in days. Growth rates were calculated by changes in tumor fluorescence area over time (µm^2^/day). Blue and green dotted circles designated to highlight relatively early growth burst of GPC6 tumors follow but rest while 3xCr tumor growth bursts later in survival. E) Representative images from two-photon imaging at different depths of tumor cells (cyan) and neurons (magenta). Visualized depth labeled on left. Age at visualization labeled above images. D) Long-term optical stability of cranial windows: widefield (top) and 2-p (bottom) images from the same animal at P62, P127, and P189. Scale (top) = 1 mm, scale (bottom) = 0.1 mm E) Examples of dual-indicator widefield images of different tumor fluorescence and activity indicators over time. Top row: thy1-GCaMP6s line and tumor pseudocolored in magenta and cyan respectively. Bottom row: iGluSnfr and tumor pseudocolored in yellow and cyan, respectively. Scale = 1 mm

### Cortical excitability and glutamate dynamics are uncoupled during GBM progression, depend on tumor genotype

Given the specific increase in excitatory, glutamatergic synapses in GPC6 tumors, we sought to investigate whether the glutamate release would be affected and what effect this would have in network activity between 3xCR tumors and those expressing GPC6. To this end, we virally introduced genetically encoded fluorescence indicators for calcium (thy1-GCaMP6s) and glutamate (AAV-FLEX-iGlusnfr + AAV-Ef1α-Cre). In 3xCr animals, we observed a rapid and spatially heterogeneous elevation of calcium activity taking place during the early period (P56-P69), followed by an indolent period with little further change (<10% between time points) in baseline calcium (Fig 4A). This suggests a low impact of the invading tumor cells on baseline neuronal calcium levels distant from the 3xCR tumor over a 6-week period. In contrast to the neuronal calcium signals, there is a steadily changing glutamate signal between similar time points (Fig 4B). At early stages, glutamate begins to accumulate before the apparent arrival of dense cortical tumor signal (P59-P63), then intensifies and extends far beyond the labeled tumor margin. As the tumor progresses, the glutamate signal engulfs the entire tumor. Also striking is the continued dynamic instability (regions of dark red and blue in colored images corresponding to large CV fluctuations) of glutamate levels throughout the growth period.

**Figure 4:**
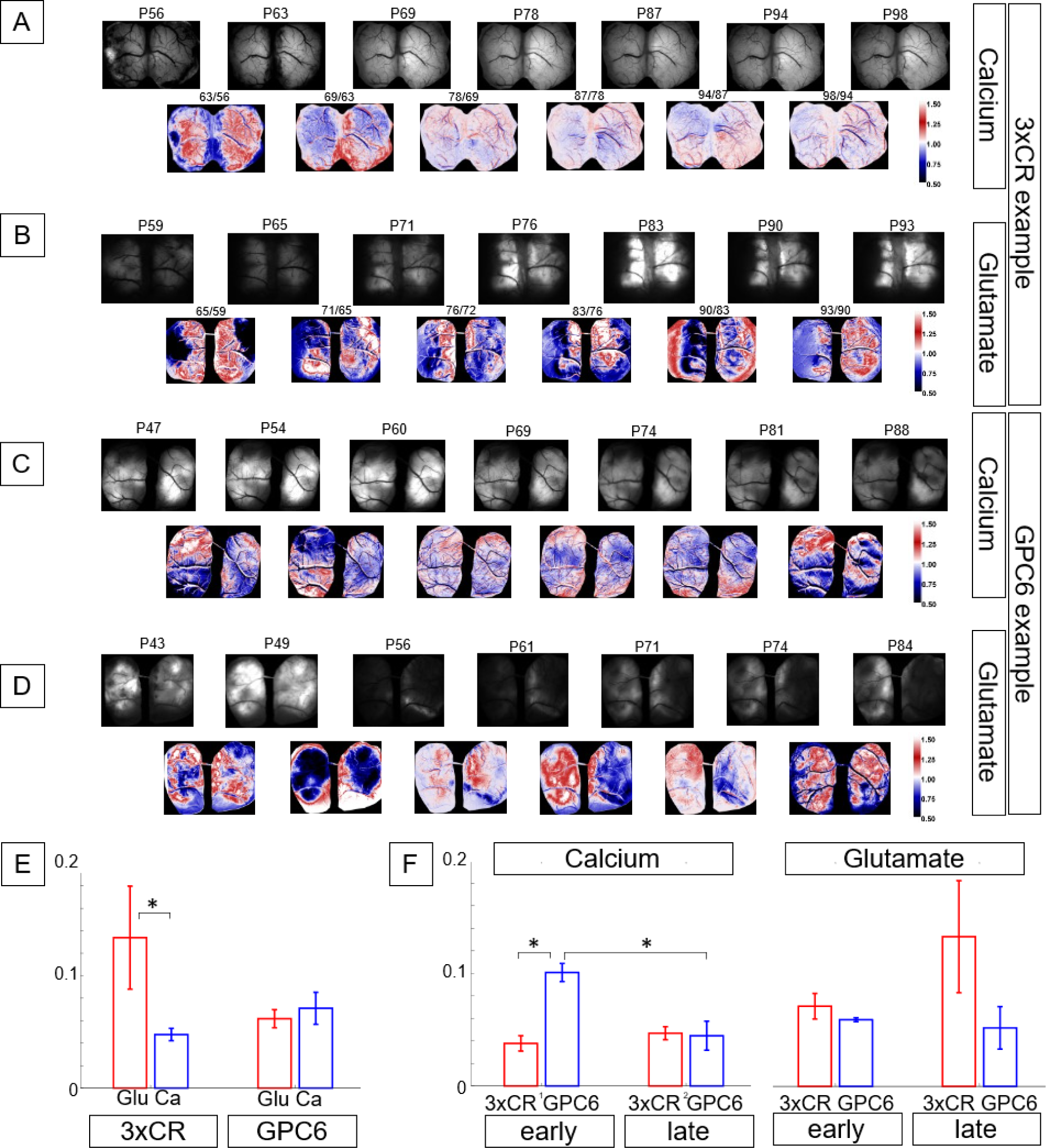
Abnormal glutamate accumulation outpaces spatial extent and temporal increase in baseline neuronal calcium activity in 3xCR tumor cortex. GPC6 baseline calcium signal is elevated at early time points (<P70) A) Top: Black/white panels show calcium fluorescence of the widefield FOV in a 3xCR tumor cortex at 7 time points between P56 and P98; equal brightness scale in each image. Bottom: Colored panels show the rate of change in tumor fluorescence at weekly intervals, generated by dividing two B/W images from consecutive timepoints, normalizing by the number of days between those sessions and by the mean of the resulting image, applying a colormap (imageJ, “union jack”) and setting the scale limits to [0.5 1.5]. B) Analogous to A), B/W panels show the evolution of baseline glutamate fluorescence intensity in a different 3xCR tumor animal between P59 and P93 with equal brightness scaling. Note the distinct nonlinear, progressive changes in signal intensity not seen in the calcium reporter mouse (A). Colored panels were constructed and scaled as in A). C) Analogous to A), B/W panels show calcium baseline signal snapshots from a GPC6 tumor animal between P47 and P88, as well as the CV/day fluorescence change between time points, revealing high early calcium signal that subsides gradually over time. D) Analogous to B), GPC6 peritumoral glutamate panels show nonlinear signal evolution over time. E) To capture the dynamical changes in fluorescence between time points, we computed the CV (SD/mean), normalized by the number of days between recordings. To account for potential sampling bias/undersampling, we extrapolated a 90^th^ percentile datapoint for each animal instead of using the maximum value. Data shown from 10 3xCR calcium-reporter mice, 5 3xCR glutamate reporter mice, 8 GPC6 calcium-reporter mice, and 2 GPC6 glutamate reporter mice. In 3xCR animals, glutamate CV/day reached 64% higher values than calcium across all time points (mean 0.134±0.046 sem vs. 0.048±0.0055 sem, p=0.01, Wilcoxon ranksum test). By contrast, GPC6 did not show a significant difference between glutamate and calcium baseline taking all time points into account (GPC6: mean glutamate = 0.062 ±0.008 sem vs. calcium 0.071±0.014 sem, p=0.6, WR test). F) Comparison of calcium and glutamate signals between early and late time periods in 3xCR vs GPC6 tumors: GPC6 calcium baseline at early time points (<P70) was higher than 3xCR early, and both GPC6 and 3xCR late (P>70): mean GPC6 calcium early/late: 0.1 ±0.008 sem / 0.047±0.006 sem, mean 3xCR calcium early/late: 0.038 ±0.007 sem / 0.045 ±0.013 sem, p = 0.0082, KW/mc test). Glutamate levels were not significantly different between the two genotypes and time periods.

In GPC6 tumors, there was a stronger change in calcium signal at earlier time points (Fig 4C), and glutamate dynamics were less intensely modulated at late time points (Fig 4D) compared to 3xCR. More specifically, across the duration of the experiment, changes in calcium baseline fluorescence appeared consistently smaller than the changes in glutamate reporter brightness in 3xCR tumor animals, but not in GPC6 (Fig 4E). However, GPC6 calcium signals accumulated at earlier time points (<P70, Fig 4F, calcium, early, blue vs red), whereas they did not change over time around 3xCR tumors (Fig 4F, calcium, 3xCr, early vs late).

This result demonstrates a significant dynamic change in extracellular glutamate levels during the progression of GBM invasion that - in non-tumor regions - correlates poorly with neuronal calcium activity levels. These distant nonlinear glutamate accumulations may precede intimate contact with tumor cells and extend several millimeters away from the tumor margin, even including large areas remote from the tumor in the contralateral hemisphere. Increased synaptogenesis in GPC6 tumors appears to significantly change the time course of these neurotransmitter modulations.

### Neural activity patterns are significantly elevated during periods of accelerated tumor growth

Specific types of GBM can display nonlinear proliferation patterns over time^9^, and this might influence their effects on surrounding neuronal physiology in their microenvironment. We asked whether spontaneous peritumoral aggregate neural activity within <7.5 mm of the tumor edge was significantly modulated with the advent of tumor cell invasion. Specifically, we analyzed whether changes in ongoing activity levels and patterns depended on concurrent tumor growth rates. To investigate this, we bifurcated the data into two bins of fast and slow growing tumors at above or below 10^5^um^2^/day, respectively (Fig 3D). When we quantify all the measured events in a given frame during fast and slow growing stages in our 3xCr brain (Fig 5A), we observed 13% fewer events/sec. This difference is absent in GPC6 brains (Fig 5B).

**Figure 5:**
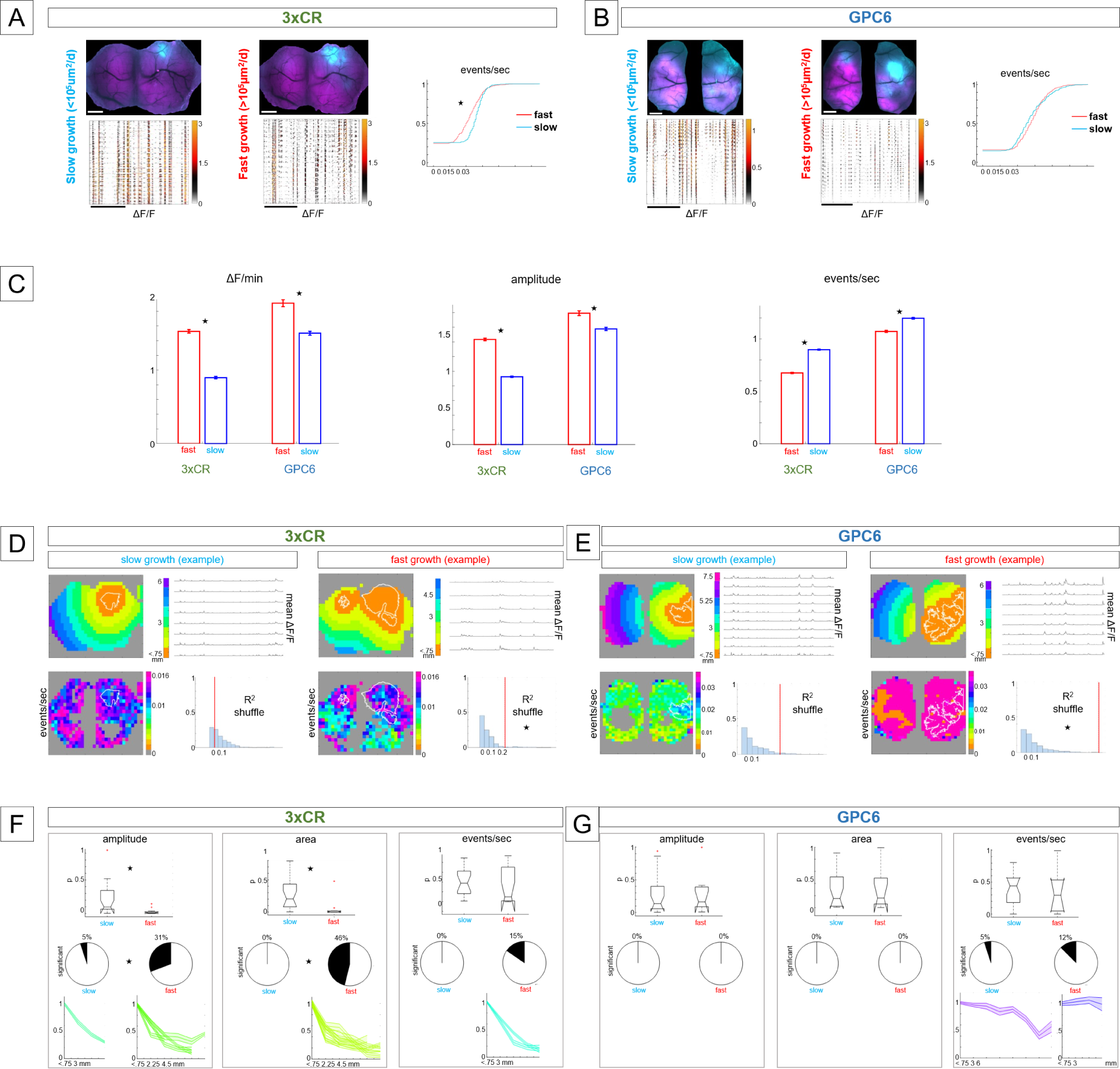
Mesoscopic neural activity patterns are significantly elevated when the tumor growth rate exceeds 10^5^µm^2^/day; Neural activity patterns drop off with distance from 3xCR tumors when proliferation rates are high; GPC6 tumors generally do not induce this effect. **A**: Example of 3xCR whole-FOVs values for 2 time points corresponding to slow (3.3*10^4^ µm^2^/day) tumor growth rate (left) and fast (2*10^5^ µm^2^/day) tumor growth rate (middle). Top panels: tumor (cyan) and calcium (magenta) FOV of the same animal at different time points. Horizontal scale = 1mm. Bottom: representative filtered and denoised ongoing calcium traces, one row per pixel (after spatial downsampling, 520 pixels total, color scale: 0 to 3 x 10^-3^ ΔF/F).). Scale = 50 sec. Right: exemplary cumulative distribution functions (cdf’s) show identified events/sec of all pixel traces (red curve = recording during fast tumor growth, blue curve = recording during slow tumor growth). In this 3xCR example, the recording taken during a period of fast tumor growth displayed, on average, 13% fewer events/sec than during slow tumor growth (mean events/sec = 9e-4 ± 2.7e-4 sem and 1e-3 ± 2.75e-4 sem, respectively, p = 3e-6, Wilcoxon ranksum (WR/mc test) **B:** As in A, top panels show tumor (cyan) and calcium(magenta) fluorescence, and bottom panels spontaneous DF/F calcium activity of a GPC6 tumor animal from representative recordings at a slow tumor growth stage (left), and a fast growth stage (middle). Events/sec were not significantly modulated between the two time points (mean events/sec = 0.0124 ± 2.5e-4 sem, and 0.0126 ± 2.5e-5 sem, respectively. right: corresponding cdf’s). **C:** Comparative bar plots for three activity metrics (DF/min, events/sec, amplitude) including pooled data from 4 3xCR animals (20 recordings during slow tumor growth, 13 recordings during fast tumor growth) and 4 GPC6 animals (20 recordings during slow tumor growth, 8 recordings during fast tumor growth); blue bars = slow growth recordings, red bars = fast growth recordings.error bars = standard error of the mean. Star denotes significant difference between fast and slow growth conditions (P<0.001) **D:** Example of two 3xCR tumor recordings in the same animal at a time point of slow tumor growth rate (4 left panels) and fast growth rate (4 right panels). Top left panel: The distance from the tumor edge was computed for each pixel in the spatially downsampled calcium movies. Distance bands are color-coded in 0.75 mm increments. Top right: The mean ΔF/F traces to the right of each distance-band plot depict a 30-sec period of mean quiet spontaneous activity corresponding to the adjacent color scale. Bottom left: To illustrate the approach, each pixel was color-coded according to the mean event rate across the analyzed recording time. Bottom left: neuronal event rates were fitted against distance from the tumor to generate a goodness-of-fit R^2^ value. Pixel distances were circularly shuffled 500k times to generate a null distribution of the linear fit R^2^ (blue histogram) and to compute the corresponding p-value of the fit significance. (red line in the histogram of the shuffled null distribution, p = 0.62). In the example from a fast-growth epoch (same animal) at the right, event rates were significantly higher close to the tumor than farther away (p = 0.02)). **E:** As in A, here we show example data from a GPC6 tumor animal from a recording during slow tumor growth (4 left panels), and one while growing fast (4 right panels): color-coded distance bands around the tumor, 30 seconds of spontaneous activity corresponding to those distances, color-coded event rate levels for all pixels, and significance of linear goodness-of-fit R^2^ with shuffled null distributions. **F:** 3xCR tumor animal recordings with significant relationships between distance and activity pattern metrics were identified using the method shown in A and B. For each metric, boxplots contain the corresponding R^2^ p-values from 20 recordings acquired during slow tumor growth (left), and from 13 recordings during fast growth (right). The horizontal line corresponds to the median, the vertical extent of the box equals the interquartile range (25th to 75th percentile), the whiskers extend to the most extreme data points not considered outliers, and the outliers are plotted individually using the ’+’ marker symbol. Notches display the variability of the median between samples, and boxes whose notches do not overlap have different medians (at α = 0.05). Pie charts show the percentage of significant recordings under both growth conditions for each metric. The shaded errorbar plots (mean and sem) underneath contain metric data from the corresponding significant recordings, normalized by the first value (corresponding to <0.75 mm distance from the tumor edge) of each recording. The x-axes of these plots extend out to variable maximal values, depending on the tumor coverage of each FOV constraining space for the distance bands. **G:** GPC6 recordings (n=4 animals) were analyzed as in C. Data are from 20 recordings under slow growth conditions and 8 recordings during fast growth. Note the lack of significant relationships between distance and activity metrics in contrast with the 3xCR recordings.

To gain a more granular view of activity level and pattern differences between fast and slow growing phases, we extracted and analyzed individual calcium events corresponding to simultaneous electrical activity of local neuronal ensembles^34–36^, and observed stark differences between the 2 stages. The differences in neural activity patterns between slow and fast tumor growth phases were highly pronounced in 3xCR animals (figure 5C). The mean activity (ΔF/min) was 70% greater during fast growth (1.5 ± 0.024 sem vs. 0.9 ± 0.016 sem, respectively, p = 7e-8, WR/mc test), indicating a strong intensification of the overall synaptic and somatic activity. The amplitudes of isolated calcium events were higher by 55% (1.43 ± 0.017 sem vs. 0.924 ± 0.008 sem, p = 1.6e-20, WR/mc test). On the other hand, the mean event rates were 25% lower when tumor growth rates were high (0.675 ± 0.006 sem vs. 0.9 ± 0.005 sem, p = 2e-170, WR/mc test). Consequently, peritumoral aggregate activity patterns consisted of less frequent but significantly larger bursts of activity when 3xCr tumors grew rapidly than during periods of slower growth.

By contrast, neural activity patterns in GPC6 tumor animals were higher overall than their 3xCR counterparts but exhibited smaller differences between periods of fast and slow tumor growth (figure 5C): The mean ΔF/min values were 27% higher (1.91 ± 0.05 sem vs. 1.5 ± 0.026 sem, p = 5e-6, WR/mc test) when tumors grew fast, mean amplitudes were 13.6% higher (1.79 ± 0.03 sem vs. 1.58 ± 0.02 sem, p = 1.5e-18, WR/mc test), and events/sec were 10.5% rarer (1.07 ± 0.01 sem vs. 1.2 ± 0.008 sem, p = 2e-170, WR/mc test). Thus, it appears that the difference in neural activity surrounding fast and slow growing GPC6 tumors was less pronounced than surrounding 3xCR tumors: in GPC6 animals, calcium events were 10.5% less frequent, versus 25%, and maximum firing rate (expressed in the transient amplitude) was 13.5% higher, whereas it was 55% in 3xCR animals.

### Tumor neuronal crosstalk depends on GPC6, tumor distance, and growth rate

Having established a differential influence of GPC6 on global neuronal activity changes and growth rate, we next asked whether activation of this cross-talk also depended on distance from tumor cells. We calculated distance bands in 0.75 mm increments, as shown in figure 5D and E for 3xCR and GPC6, respectively, averaging the ΔF/F signal concentrically from the tumor border (outlined in white lines). We determined significant differences by comparing event rates to a R^2^ goodness of fit value analysis. In both 3xCr and GPC6 tumor brains, there was a significantly higher event rate proximal to the tumor (p-values .02 and 2e-6, respectively) during the fast growing phase. Interestingly, the R^2^ of event rate as a linear function of distance from the tumor during slow tumor growth was not significant in either tumor model. This suggests that distance is only a relevant factor underlying cross-talk during the fast growth phase.

Subsequently, we applied this analysis to the activity metrics (amplitude, area and event rate (“ΔF/min”, and “rhythmicity” are shown in suppl figure 3): In 3xCR animals, linear fit p-values (distance from tumor edge vs. activity metrics) were significantly different between slow and fast tumor growth periods for event amplitude and area, but not event rate (Fig 5F). To analyze whether the number of significant recordings differed between the two growth rate conditions, we performed a χ^2^ test between the percentage of recordings that had a significant linear relationship between distance and activity metrics (pie charts). In 3xCR animals, calcium event sizes were significantly different depending on tumor distance: mean amplitudes were more likely to drop off with distance when growth was fast rather than slow (mean p-values: 0.22 ± 0.05 sem and 0.0273 ± 0.01 sem respectively, p = 0.001, WR/mc test). We determined that 31% of recordings in fast tumor growth states were significant vs. 5% during slow growth (p = 0.04, χ^2^ test). Mean amplitudes in slow growth recordings were 70% lower at 3mm from the edge, and 80% lower at 3.75mm under fast growth conditions. Similarly, calcium event areas had a stronger association with distance under fast than slow growth (mean p-values: 0.047 ± 0.05 sem vs 0.28 ± 0.05 sem respectively, p = 0.002, WR test), and 46% of recordings were significant vs 0% (p = 8e-4, χ^2^ test). Mean area values were on average 83% lower at a distance of 4.5 mm than <0.75mm from the tumor margin. However, event rates were, in most recordings, not significantly associated with distance from the tumor, and there was no significant difference between slow and fast growth periods (mean p-values: 0.4 ± 0.06 sem vs. 0.34 ± 0.105 sem, p = 0.32 WR test). While 2 recordings during fast growth showed significance, there were none during slow growth, but this difference was not significant (p=0.07, χ^2^ test). When significantly modulated, event rate was on average 83% lower at a distance of 3.75 mm from the tumor edge.

Whereas these results from 3xCR tumor animals indicate that overall calcium events were progressively diminished at increasing distances from the tumor, GPC6 peritumoral activity generally did not follow the same activity-distance relationships (Figure 5G). There were no recordings with a significant correlation of distance and amplitude or area (empty pie charts in fig 5G, see suppl fig 3 for ΔF/min, rhythmicity), and goodness-of-fit p-values were never different significantly different between recordings from slow and fast growing tumor epochs (amplitude: 0.28 ± 0.07 sem and 0.29 ± 0.11 sem, p = 0.65 WR test; area: 0.27 ± 0.07 sem and 0.3 ± 0.13 sem, p = 0.8, WR test). However, the event rate was significantly modulated as a function of distance in two GPC6 recordings, one while the tumor was growing slowly, and one fast. In accordance with these low numbers, there was no significant difference between the two conditions (mean p-values: 0.38 ± 0.06 sem and 0.34 ± 0.12 sem, p = 0.7 WR/mc test), nor between the percentage of significant recordings (5% vs 12%, p = 0.49, χ^2^ test). In the significant slow growth recording, mean event rates were 16% lower at a distance of 3 mm, and 42% lower at 6 mm. The maximum distance in the recording taken during a period of fast tumor growth was 3 mm.

These results demonstrate that mesoscopic peritumoral neural activity patterns can be differentially modulated according to distance from the tumor margin at a sub-millimeter scale, depending on the contemporaneous proliferative state and specific genetic makeup of the tumor. Calcium event amplitude and area were generally stronger close to 3xCR tumors under rapid growth, even at distances of 2-3 mm. GPC6 tumors did not seem to influence calcium event strength in this distance-dependent manner, but we did observe a modest but significant distance dependent difference in event rate.

### GPC6 dependent changes in cellular level activity patterns inside versus outside the tumor margin

One-photon widefield imaging enabled monitoring of large cortical areas simultaneously, proving useful to measure activity changes around tumors with a circumference of several mm. However, it does not permit cellular resolution recordings, and the mesoscale signal is dominated by dendritic and axonal activity that may obscure the underlying soma voltage signal^37, 38^. Therefore, we used awake 2-photon high resolution calcium imaging to analyze somatic neuronal activity patterns and ensemble metrics at the cellular level in 12 3xCR, 9 GPC6 and 8 non-tumor control animals (suppl fig. 4) over time. First, we asked whether activity patterns were disrupted between intra- and extramarginal areas (see figure 6A for example recordings, 6D for neuron-tumor images). In 3xCR tumor animals, intramarginal event rate was elevated by 22% compared to extramarginal (>1 mm) FOV’s (p = 6e-4, WR/mc test), indicating a shift in firing patterns towards shorter, more frequent bursts. (figure 6B, left). On the other hand, neurons inside GPC6 tumor margins were substantially more active overall than those in the periphery (+71.5%, p=1e-9, WR/mc test), and their event amplitudes were higher by 54.4% (p=1e-23, WR/mc test, figure 6B, right).

**Figure 6:**
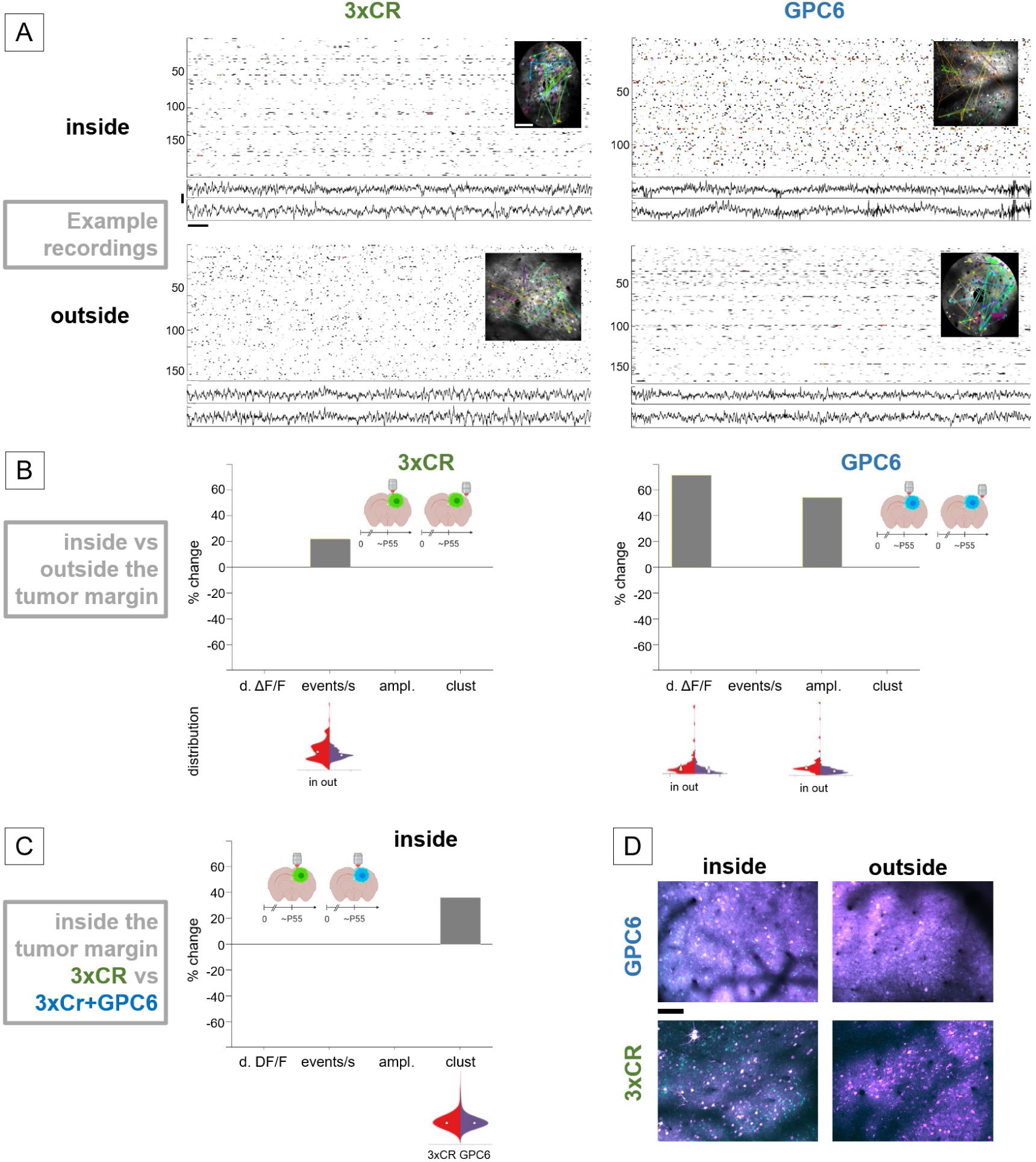
3xCR and GPC6 alter neuronal activity inside and outside the tumor margins in distinct ways. a) Comparison of calcium activity inside tumor margins (top two panels) vs outside (bottom two panels), for 3xCR tumors (left) and GPC6 tumors (right). Each panel consists of a raster plot of deconvolved calcium activity (one row per neuron), 2-ch EEG (vertical scale = 200µV), and an insert showing the FOV with clustered neurons in matching colors and line connecting pairs of neurons within the clusters (line width proportional to connection strength). The “3xCR inside” and “GPC6 outside” recordings were acquired in spiral scan mode, whereas the “3xCR outside” and “GPC6 inside” FOV’s were imaged under resonant scan. Horizontal scale = 1 sec. b) Comparison of activity metrics between neurons inside and outside the tumor margin for 3xCR (left) and GPC6 (right) tumor animals. Only significant changes for mean deconvolved ΔF/F, mean events/sec, mean event amplitude, and mean clustering coefficients are shown in the bar graphs. Below the bar graph are violin plots of the distributions whose comparison is depicted above. The brain section pictogram inserts indicate tumor type (3xCR=green, GPC6=blue), time bins, and imaging locations. c) Comparison of activity parameters recorded inside Cr86 tumor margins versus GPC6 tumor margins. d) Example of GPC6 FOV’s inside and outside the tumor margin (top), analogous for Cr86 tumors (bottom). Tumor cells = cyan, neurons = magenta. Scale bar = 100µm

Unexpectedly, this result showed a significant difference in somatic activity patterns inside vs. outside the tumor margin, with strong dependence on tumor genotype, and with a stronger drop-off with distance in GPC6 animals, not 3xCR. However, the datasets used here were taken at earlier timepoints than the widefield imaging data (Fig 5): 3xCR inside: mean age (day P) = 57.6 ± 4.8 SD, GPC6 inside: 49.8 ± 7.9 SD, outside 3xCR: 54.2 ± 2.5 SD, outside GPC6: 53.8 ± 2.3 SD, control: 53 ± 6.1 SD. The previous mesoscopic analysis (Fig 5) taken during periods of rapid growth were at a mean age of 96.3 ± 23 SD (3xCR), and 77 ± 17.5 SD (GPC6), on average at least 20 days later. This agrees with earlier observations of 3xCR tumors showing more enduring activity than their GPC6 counterparts (Fig 3).

A direct comparison of activity inside 3xCR and GPC6 tumor margins demonstrated that dΔF/F activity production, event rates, and amplitudes were not significantly different between 3xCR and GPC6 tumors. However, GPC6 CCs were 36% higher than 3xCR (p = 0.032, WR/mc test, figure 6C). (GPC6 clustering was 26% higher than controls but not significant, and 3xCR was 7% lower than controls but not significant). Together, these results suggest that both 3xCR and GPC6 tumors alter the activity patterns of neurons inside their margins with respect to more distant populations. Specifically, 3xCR tumors favored firing patterns composed of frequent and shorter bursts, and GPC6 tumors caused overall less activity beyond the tumor margin, but higher instantaneous firing rates within the tumor margin.

### Dynamic changes in neurons outside the leading edge reflect GPC6 expression

To probe the range of tumor influence analogously to the growth dynamics analysis (Fig 3), we asked whether neurons located 1-2 mm beyond the tumor margins are affected differentially over time depending on tumor genotype (Fig 7). In 3xCR tumor animals, cellular mean dΔF/F activity was 6.5% lower at mid vs early time points (p=6e-4, WR/mc test, figure 7B), however event rate was 44% higher (p=1.6e-29, WR/mc test), indicating disorganized, less burst-dominated activity patterns with much shorter events. Mean amplitude was 7.4% higher at mid time points (p=0.002, WR/mc test). Mean CCs were reduced by 70% (p=2e-15, WR/mc test), further pointing towards an early dysregulation of local functional ensemble structure. From mid- to late time periods, there was a small increase in mean dΔF/F activity levels (+4.3%, p = 0.0014, WR/mc test), and a small reduction in event amplitude (-8.2%, p = 1.8e-4, WR/mc test).

**Figure 7:**
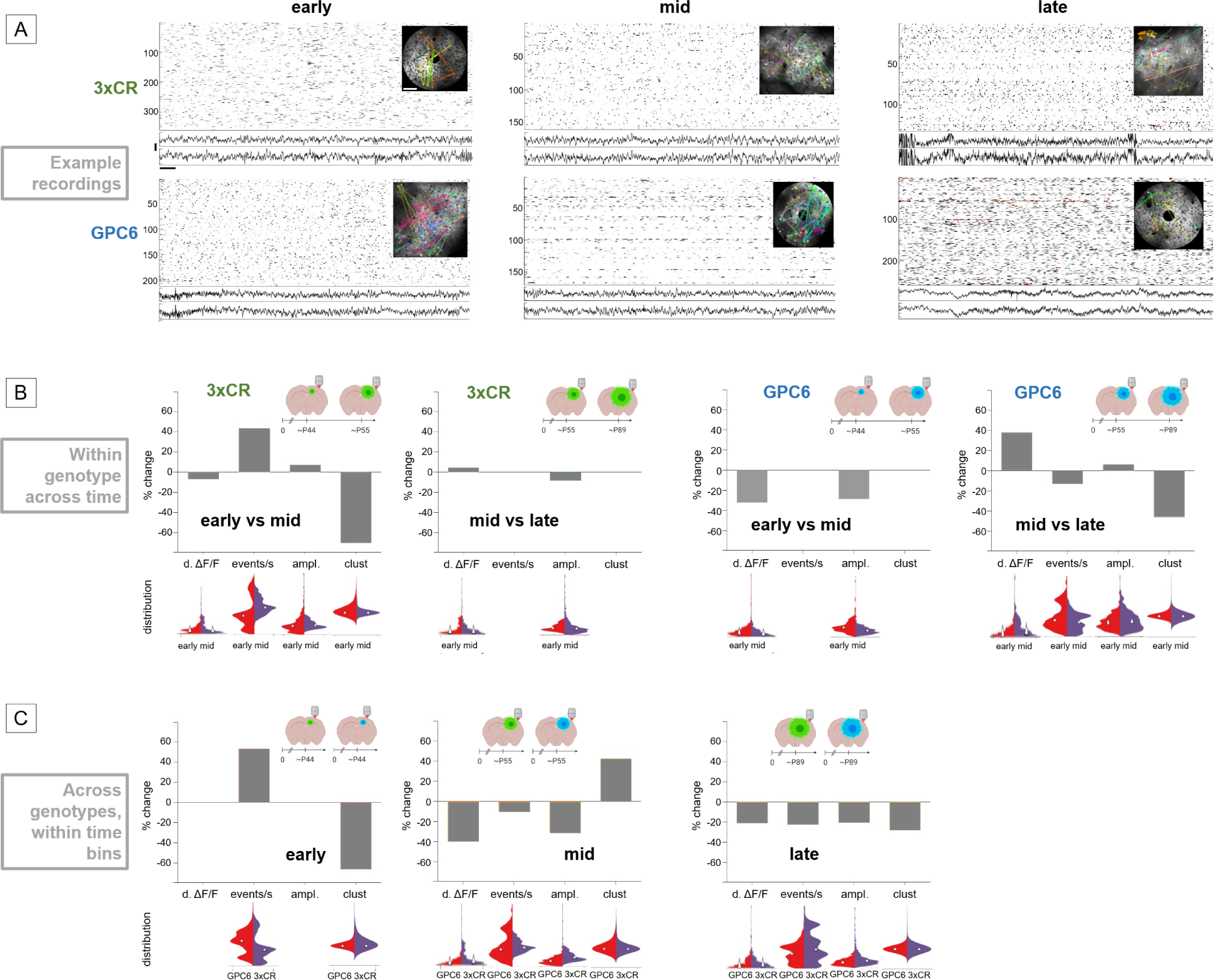
Distinct temporal dynamics of 3xCR and GPC6-induced changes in neuronal activity patterns located 1-2mm beyond the tumor margin. A) Example plots of dΔF/F, EEG and clustering for early (P41 - 49, mid (P49-56 and late (P68-129) time bins, analogous to figure 6A: dΔF/F, EEG’s (vertical scale = 200 μV, horizontal scale = 1 sec), and FOV insert with neuronal clusters (white scale bar = 100 μm. “3xCR early”, “GPC6 mid” and “GPC6 late” examples were acquired in spiral scan mode, the rest in resonant scan mode. B) Comparison of activity metrics across 3 time bins for both tumor genotypes: Analogous to figure 6, each bar plot shows (from left to right) significant percent changes in neuronal deconvolved dΔF/F activity, mean event rates, mean calcium transient amplitudes, and clustering coefficients across time points for both tumor genotypes separately. The two left panels follow changes in 3xCR tumor-induced activity patterns from early to mid time bins, and from mid to late time bins. Violin plots of distributions whose comparisons yielded significant differences are shown underneath. The two right panels show changes in the same parameters and time periods for GPC6 tumor animals. C) Here we compare the activity patterns directly between GPC6 and 3xCR tumors, separately for 3 time bins: early (left), mid (middle), and late (right).

In GPC6 animals, cellular dΔF/F activity was 32% lower at mid time points compared to early (p = 8e-9, WR/mc test), and mean amplitude was reduced by 29% (p = 2.4e-21, WR/mc test), indicating lower firing rates inside otherwise unchanged bursts, and no significant change in functional synchrony. By contrast, from mid to late time points, mean dΔF/F activity increased by 38% (p = 1e-4, WR/mc test), mean rate of Ca events/sec was 13% lower (p=5e-8, WR/mc test), event amplitudes increased by 6% (p = 0.036, WR/mc test), and mean CC decreased by 46% (p = 1e-8, WR/mc test).

These results indicate that both tumor types alter extramarginal neural network activity over time but with different temporal dynamics. Specifically, 3xCR caused significant disorganization of activity patterns early on, whereas GPC6 tumors moderately reduced firing and intra-burst frequency early, but later caused a large dΔF/min increase with a reduction in functional organization.

Lastly, we directly compared the two tumor genotypes within the 3 time bins to quantify differences in their impact on extramarginal neuronal activity parameters during progressive stages of tumor invasion (figure 7C). At early time points, neurons around GPC6 had a higher mean event rate (53% increase, p = 8.9e-12) than around 3xCR tumors, and their mean CC was 66% lower (p = 3.2e-6, WR/mc test). At mid time points, mean dΔF/F was 40% lower around GPC6 tumors than around 3xCR tumors, (p = 3.2e-26, WR/mc test), event rate was 10% lower (p = 0.004, WR/mc test), mean amplitudes were 31% lower (p = 1e-53, WR/mc test), but CCs were 42.7% higher (p = 0.003, WR/mc test), suggesting overall less activity but stronger functional organization within neuronal populations around GPC6 tumors. At late time periods, dΔF/F, event rates and amplitudes were further reduced by 20.5% (p = 0.007, WR/mc test), 22% (p = 1.9e-11, WR/mc test) and 20% (p = 1.4e-13, WR/mc test), respectively. CC was 28% lower (p = 1.4e-7, WR/mc test). Taken together, differences between GPC6 and 3xCR peritumoral neuronal patterns underwent several transitions, from more frequent but shorter and disorganized bursts in GPC6 tumors at the early stage, to less activity with increased organization at the mid stage, followed by overall lower activity patterns and organization at late time points compared with 3xCR tumors.

## Discussion

For many decades, GBM has consistently ranked as one of the least successfully treated human malignancies, with the highest recurrence and lowest survival rates, despite the introduction of promising therapeutic interventions ^19^. In addition to poor survival prognoses, GBM routinely occurs with severe neurological comorbidities including early seizure onset and cognitive deficits ^2, 39^, suggesting a functional interrelationship. A spectrum of mechanisms underlying this hyperexcitability ^20, 40–43^ has been proposed, supported by several studies indicating that anti-epileptic drugs may slow tumor progression^1, 44–46^.

Despite these advances, many questions about the reciprocal feedback mechanisms between GBM and the surrounding neural activity and their spatial extent remain unanswered. Here we use a recently established GBM model of native glioma growth in an immunocompetent host ^4, 17^, a model that recapitulates many hallmarks of the human disease^7^, to chronically visualize and correlate glutamate, neuronal hyperactivity, and tumor expansion in vivo at high temporal resolution for the first time. We show evidence of early extensive cortical glutamate accumulation as 3xCR tumor cells invade cortex that is not accompanied by equally dramatic changes in baseline calcium signaling, suggesting that it does not depend exclusively on synaptic release, supporting ample evidence of multiple molecular pathways that can alter the GBM extracellular microenvironment by raising free glutamate levels ^21, 43, 47–49^.

GBM growth rates vary in human, and we replicated this feature in our finding that tumor growth dynamics were not sustained into late stages (as seen in the 3xCR model, Fig 3) when we added overexpression of a single glypican gene linked to synaptogenesis (GPC6) to the IUE CRISPR construct. Likewise, we saw an early surge in baseline aggregate (mesoscopic) calcium signal not seen in 3xCR animals (Fig 4). Furthermore, we established that distinct cellular neuronal network activity disruption, both between intramarginal and extramarginal neuronal populations, and over time at locations >1mm beyond the tumor edge, depend on the genetic makeup of the tumor. Recent work has implicated several subtypes of glypicans, astrocytic secreted proteins that promote glutamate receptor expression and induce functional synapse formation ^23, 50^. We previously showed that *GPC3*, a member of the glypican family, can drive increased peritumoral neo-synaptogenesis when expressed in GBM using our IUE CRISPR/Cas9 model ^17^. The GPC6 result was somewhat unexpected, since like GPC3, GPC6 tumors also promote peritumoral neosynaptogenesis (Fig 2G-H). Further analysis will be needed to understand the disparate contributions of GPC homologs to tumorigenesis.

Analyzing the mesoscopic calcium activity patterns in both 3xCR and GPC6 animals revealed that multiple activity metrics were significantly shifted when tumor growth rates were high versus periods of slow tumor growth, favoring fewer but stronger calcium events. However, GPC6 tumors had a less pronounced relationship between growth rate and activity patterns. At a finer spatial scale, we found that 3xCR tumor animals showed a stronger difference in calcium activity profiles between periods of fast and slow tumor growth: calcium events were larger close to the tumor edge when the tumor was growing fast, whereas in GPC6 recordings we only saw sporadic hints of a distance relationship with event rate. These results indicate an unexplained but important difference in the way specific genetic makeup of GBM may influence how it interacts with the surrounding microenvironment.

3xCR tumors caused early reduction in somatic neural activity clustering with shorter more frequent bursts compared to GPC6 recordings, while at mid points they favored lower overall activity and firing rates, and at the late stage all activity metrics were lower than those around GPC6 tumors, underscoring the significant differences in tumor/neuron crosstalk at the leading edge. Our current understanding that tumor genetics drive progression of the cortical hyperexcitability microenvironment and hence impairment of higher order cortical function predicts that matching precision GBM diagnoses with information on residual disease can help guide future tailored management of the neurological comorbidity of brain tumors.

## Acknowledgements

The authors wish to thank Ryan Ostrom and Juan Enrique Villacres Perez for technical assistance. Supported by NCI R01CA223388 (JLN, BD), R01NS124093 (BD), R50CA252125 (KY), and Blue Bird Circle Foundation (JLN).

