## Supplementary material for "Glioblastoma disrupts cortical network activity at multiple spatial and temporal scales": 4 supplementary figures + text

**Supplementary results**

Chronic calcium activity analysis at fast timescales and mesoscopic spatial scales enables analysis of the dynamical range of influence inside and beyond the tumor margin in 3xCR vs GPC6 tumor animals. Here we show a flow chart style schema of calcium widefield analysis in relation to tumor distance, and how we tested for significance between distance and activity metrics (suppl fig 1).

Results from 3xCR tumor animals indicate that calcium events were overall progressively smaller at increasing distances from the tumor - and in some recordings this was also seen for overall DF/F activity and rhythmicity (mean p-values: 0.57 ± 0.08 sem and 0.22 ± 0.09 sem slow/fast respectively, p = 0.06, WR/mc test; and mean p-values (slow/fast): 0.42 ± 0.06 sem / 0.36 ± 0.1 sem, p = 0.4 WR/mc test), percentages of significant recordings (0% vs 15%, p = 0.07, $\chi^2$ test). Rhythmicity in significant recordings taken when tumor growth was fast dropped by 83% at a distance of 3.75 mm, on average, (suppl fig 2)

There was also a small but significant increase in rhythmicity in GPC6 but not in 3xCR animals (suppl fig 3), potentially indicating a difference in the way these tumors affect functional synaptic physiology, and not just numbers, depending on their concurrent growth rates.

We examined whether intramarginal somatic activity differed in similar ways from cortical activity recorded in non-tumor control animals. We found that in 3xCR tumor animals, mean focal dΔF/F (i.e. AP production) measured inside the tumor margin was unchanged compared to controls. However, the mean event rate was elevated by 52% (p = 1e-13, WR/mc test), and amplitude by 17.7% (p = 2e-3, WR/mc test, suppl. figure 4A, left). Neuronal calcium event rates inside GPC6 tumor margins were 36% higher than controls (p = 4e-9, WR/mc test), and amplitudes 15% higher (p = 6e-5, WR/mc test). However, there was also a 26% reduction of overall calcium activity compared to controls (p = 3e-3, WR/mc test, supppl. figure 4B, right).

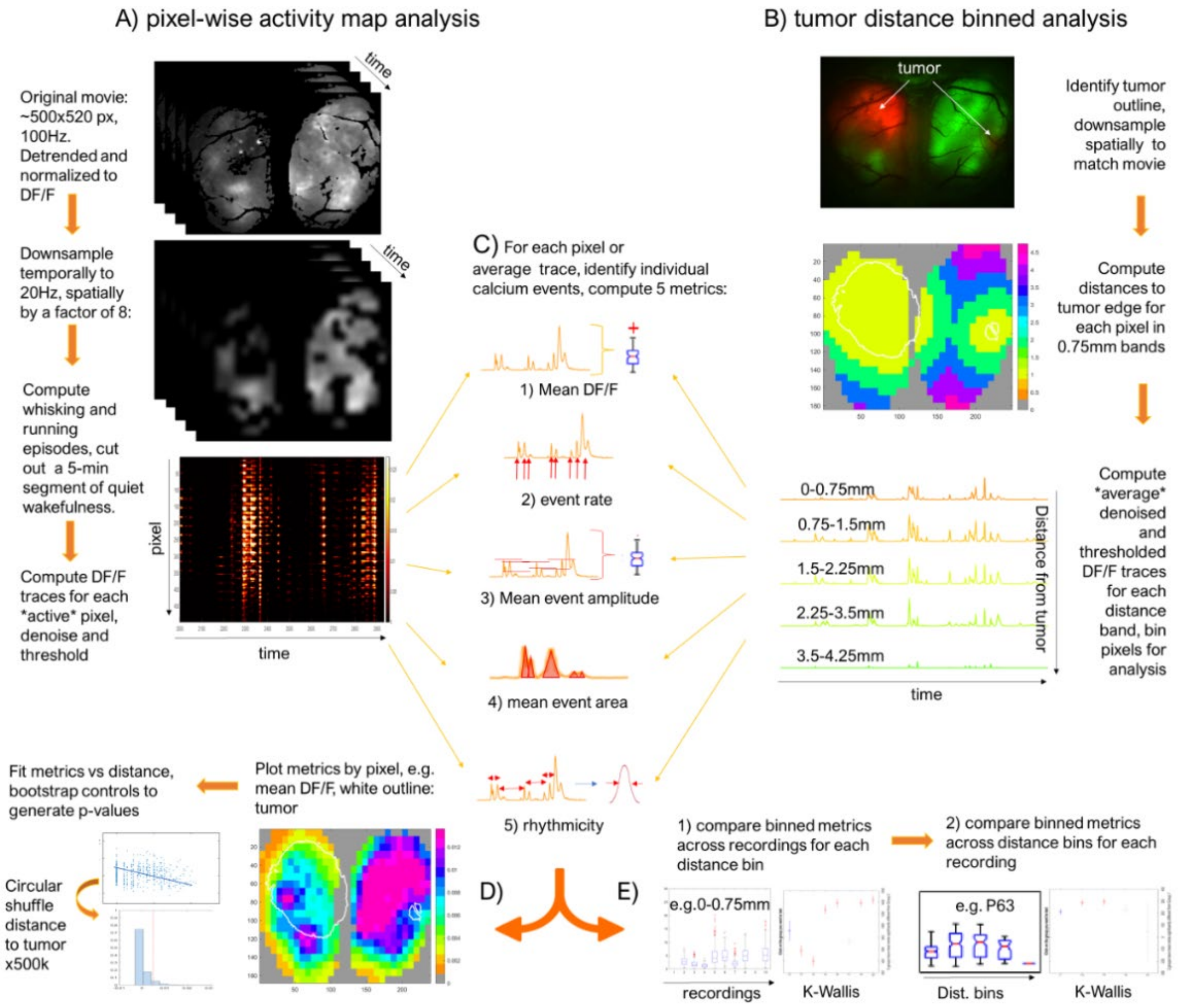


**Suppl fig 1 (supplement to Figure 1): Chronic calcium activity analysis at fast timescales and mesoscopic spatial scales enables analysis of the dynamical range of influence inside and beyond the tumor margin in 3xCR vs GPC6 tumor animals**

A) 1-p widefield images are recorded at ~500x520 pixel resolution and 100Hz sampling rate. Images are downsampled temporally to 20Hz and spatially by a factor of 8, resulting in ~0.24mm pixel resolution. After computing active whisking and running periods, 300 sec of quiet wakeful activity is selected. For each pixel, a corresponding dΔF/F signal trace is computed, detrended, and thresholded at 3 SD above mean baseline noise level.

B) Using a snapshot image taken with a green/red filter (for RFP labeled tumor) or 400nm/blue filter (for BFP labeled tumor), the tumor margin is computed for each recording at different time points. This image is aligned and spatially downsampled as in (A), and pixels are assigned to distance bands of 0.75 mm width. Average ΔF/F traces are calculated for visualization, and pixels are binned into the respective 0.75 mm distance bands.

C) Each ΔF/F trace is processed to extract the following metrics: I) mean ΔF/F over the duration of each trace, ii) calcium transient event rate $sec^{-1}$), iii) mean event amplitude for each trace, iv) mean area under the curve of identified calcium events, and v) rhythmicity, i.e. the inverse of the width at half maximum amplitude for the distribution of inter-event intervals.

D) To visualize changes in activation patterns over time, values for the computed metrics are plotted pixel by pixel with the outline of the tumor overlaid in white. To determine a possible relationship between the distance to the tumor edge and the neuronal activity metrics, the linear fit $R^2$ is computed. Next, distances to the tumor edge are scrambled via circular shuffling 500k times to create a bootstrapped null distribution and a p-value for the significance of the correlation.

E) Next, metric distributions for pixels binned into 0.75 mm-distance bands are computed, and compared, first across time for each distance bin, and then across distance for each time point (Kruskal-Wallis test with correction for multiple comparisons).

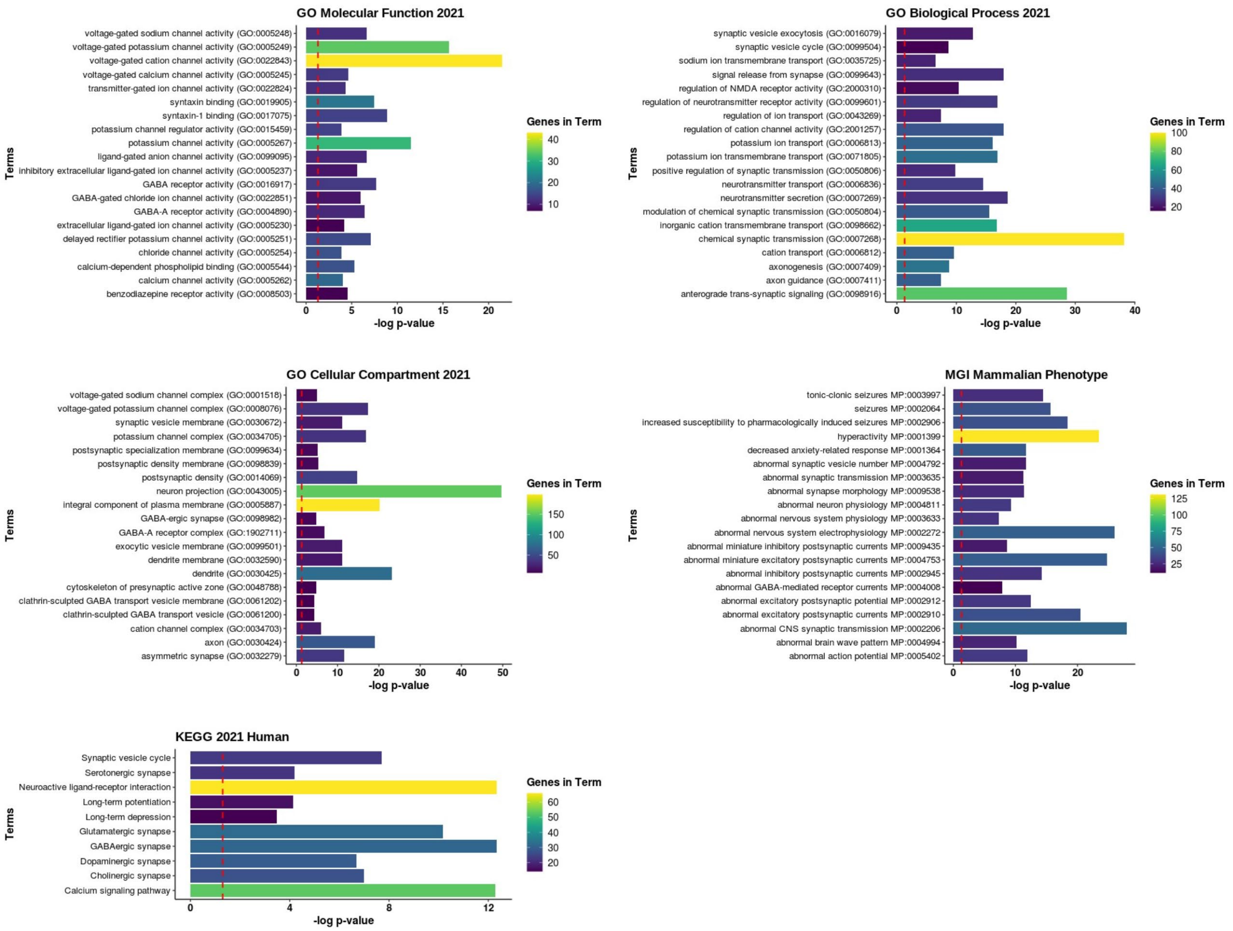


**Suppl Fig 2 (suppl to Figure 3): Bioinformatics analysis of differentially upregulated genes in GPC6 tumor brains.**

Bar graphs demonstrate of different significance values of various terms from different ontology databases. Significance values were calculated as the –log(p-value). Colors of bars indicate the number of genes in each term that were found. Dotted red line set at p = .05 threshold.

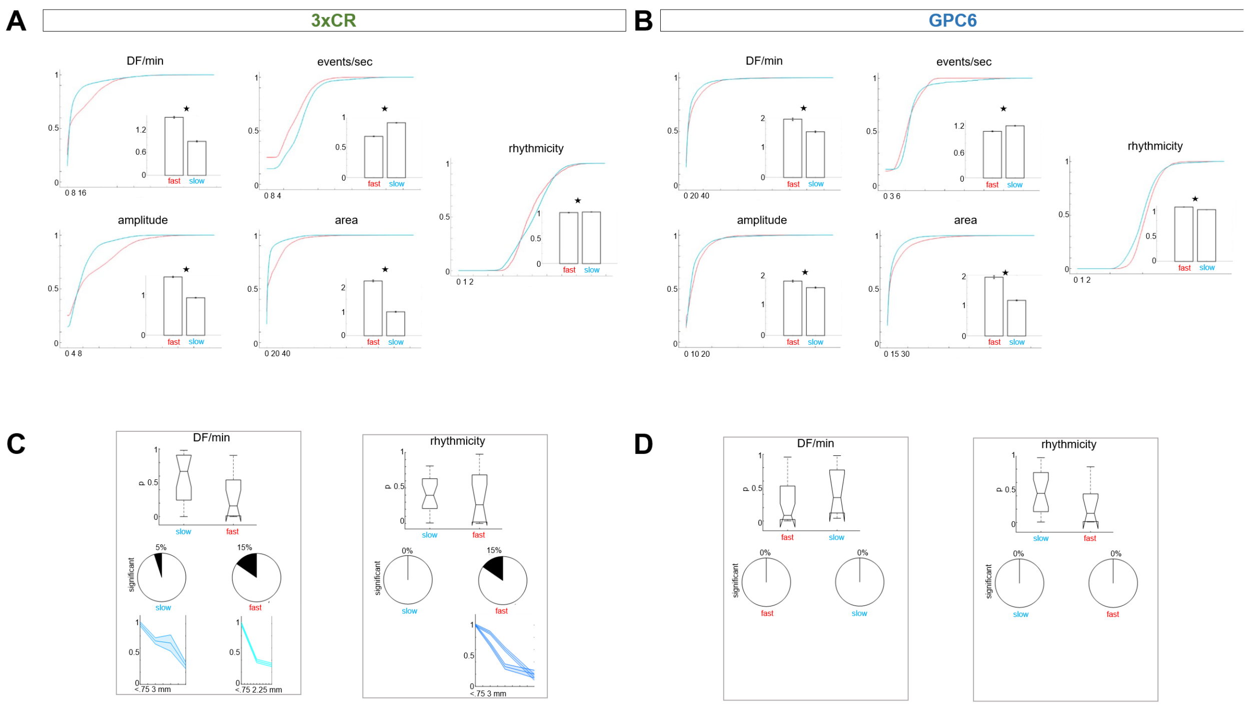


**Suppl Fig 3 (Suppl. to Figure 5): Aggregate neural activity patterns are significantly elevated when the tumor growth rate exceeds $10^5$µm$^2$/day**

**A:** Cdf plots for all five metrics (DF/min, events/sec, amplitude, area, and rhythmicity) including pooled data from 4 animals (20 recordings during slow tumor growth, 13 recordings during fast tumor growth); blue curves = slow growth recordings, red curves = fast. Each bar plot insert shows the means of the two distributions under comparison; errorbars = sem.

**B:** Analogous to C, cdf plots include data from 4 GPC6 animals (20 recordings during slow tumor growth, 8 recordings during fast tumor growth); blue curves = slow growth recordings, red curves = fast. Each bar plot insert shows the means of the two distributions under comparison; errorbars = sem.

**C:** 3xCR tumor animal recordings with significant relationships between distance and DF/min and rhythmicity metrics were identified using the method shown in A and B. For each metric, boxplots contain the corresponding $R^2$ p-values from 20 recordings acquired during slow tumor growth (left), and 13 recordings during fast growth (right). The horizontal line corresponds to the median, the vertical extent of the box equals the interquartile range (25$^{th}$ to 75$^{th}$ percentile), the whiskers extend to the most extreme data points not considered outliers, and the outliers are plotted individually using the '+' marker symbol. Notches display the variability of the median between samples, and boxes whose notches do not overlap have different medians (at $\alpha$ = 0.05). Pie charts show the percentage of significant recordings under both growth conditions for each metric. The shaded errorbar plots (mean and sem) underneath contain metric data from the corresponding significant recordings, normalized by the first value (corresponding to <0.75 mm distance) of each recording. The x-axes of these plots extend out to variable maximal values, depending on the tumor coverage of each FOV constraining space for the distance bands.

**D:** GPC6 recordings (n=4 animals) were analyzed as in C. Data from 20 recordings under slow growth conditions and 8 recordings during fast growth. Note the lack of significant relationships between distance and activity metrics in contrast with the 3xCR recordings.

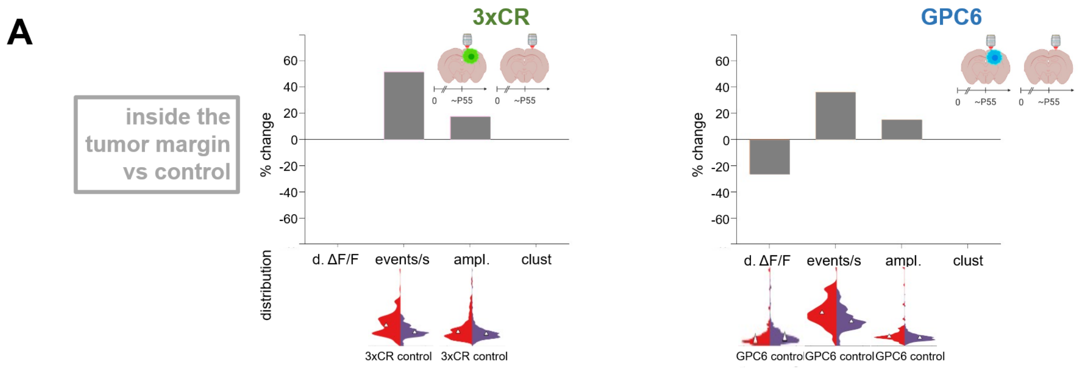


**Suppl. fig 4 (suppl to figure 6):**

A) As in fig 6 b, the left panel shows the comparison between neuronal activity parameters inside 3xCR tumor margins versus non-tumor control animals, and the right panel compares activity inside GPC6 tumor margins with controls.
